# Disturbing immune homeostasis by neutrophil loss of Uba1 induces VEXAS-like autoinflammatory disease in mice

**DOI:** 10.1101/2025.04.09.648068

**Authors:** Ge Dong, Jingjing Liu, Hongxi Zhou, Wenyan Jin, Yuchen Wen, Zhiqin Wang, Keyao Xia, Jianlin Zhang, Linxiang Ma, Yunxi Ma, Lorie Chen Cai, Qiufan Zhou, Huaquan Wang, Wei Wei, Ying Fu, Zhigang Cai

## Abstract

**Objective:** VEXAS (Vacuoles, E1 enzyme, X-linked, Autoinflammatory, Somatic) syndrome is an identified haemato-rheumatoid disease caused by somatic *UBA1* mutations in hematopoietic stem cells. We report *Uba1* loss in various mouse hematopoietic cell types leads to diverse effects, with approximately 70% Uba1 depletion in neutrophils inducing non-lethal VEXAS-like symptoms.

**Methods:** Using nine different Cre/flox-mediated conditional-knockout (CKO) models, we interrogated the phenotypes caused by hematopoietic loss of *Uba1*. Neutrophil-specific depletion of Uba1 were validated and VEXAS-like phenotypes were examined.

**Results:** *Uba1* loss in HSCs induces extensive hematopoietic cell death while in B or T cells, or megakaryocytes induces corresponsive cell death but these mutants appear normal. *Uba1* loss in monocytes and neutrophils failed to induce cell death and the mutants are viable. Among the models, only *Uba1* loss in neutrophils manifests autoinflammatory symptoms including increased counts and percentage of neutrophils, increased proinflammatory cytokines, vacuoles in myeloid cells and dermatitis. Residual Uba1 is about 30% in the mutant neutrophils, which manifest disturbed cellular hemostasis. Genetic loss of *Morrbid* partially mitigated the VEXAS-like symptoms.

**Conclusion:** Our study reveals diverse effects of *Uba1* loss in hematopoietic cells and establishes a VEXAS-like murine model, facilitating understanding and potential treatments for this syndrome prevalent in aged men.

**HIGHLIGHTS:** *WHAT IS ALREADY KNOWN ON THIS TOPIC:* VEXAS syndrome is a recently identified hematological and immunological disease prevalent in adult man but rarely in adult woman. Somatic mutations in the E1-enzyme encoding gene *UBA1* in hematopoietic stem cells is the driver on the top of the genetic etiology of the disease. However, the major pathogenic cell type(s) for VEXAS syndrome has not been experimentally examined and mouse models recapitulating the disease are lacking.

*WHAT THIS STUDY ADDS:*

i. Using nine different conditional-knockout (CKO) murine models, we interrogated the pleiotropic phenotypes caused by loss of the ubiquitin activation enzyme Uba1 in different hematopoietic cell types;
ii. Our results demonstrated that among the nine tested CKO mutants, only neutrophil loss of *Uba1* results in VEXAS-like autoinflammatory disease;
iii. The VEXAS-like symptoms in the *S100a8Cre-CKO* mutant mice include: increased counts of white blood cells and neutrophils, increased percentage of neutrophils, increased serum level of proinflammatory cytokines (IL-1β, IL-6 and TNFα), observation of vacuoles, increased survival, and increased phagocytosis in mutant neutrophils;
iv. Pharmacological treatments with IL-1 inflammatory pathway inhibitors Anakinra or Canakinumab or genetic loss of the myeloid pro-survival regulator *Morrbid* partially mitigated the VEXAS-like symptoms in the *S100a8Cre-CKO* mutants;

*HOW THIS STUDY MIGHT AFFECT RESEARCH, PRACTICE OR POLICY:* The study reports a technical strategy of developing the murine models for VEXAS syndrome. In addition, the study dissects cellular and molecular mechanisms, especially the cell-type-dependent tolerance and pathogenicity of loss of function of *Uba1*, for the occurrence of the autoinflammation diseases in mice. The study provides translational implications in etiology and potential treatment choices for the newly-identified haemato-rheumatoid syndrome in clinical management.

## INTRODUCTION

It has been recognized for almost 40 years that protein ubiquitylation, or called as ubiquitination, describing a posttranslational process about transferring the 76-amino acid peptide “ubiquitin” (∼8.5kDa) on host proteins in nucleus or in cytoplasm, is critical for nearly all aspects of eukaryotic biology and cellular homeostasis (1). Aberrant protein ubiquitylation induces cellular dysfunction in many aspects of cell activities including neuronal cells or immune cells among other important cells, and may represent signs of diseases (2). Among multiple mechanisms of ubiquitin signaling, for example, ubiquitylation of histone in nucleus regulates genome stability and gene expression (3), and ubiquitin activation in the cytoplasm is the first step of the E1-E2-E3 cascade, which are critical for almost of all of proteins inside the eukaryotic cells and maintains cell homeostasis (4). Deficient protein ubiquitylation results in inadequate protein degradation, which in turn will generate overloaded proteins inside the cell and induce unfolded protein stress response (UPR) (5, 6).

In the E1-E2-E3 cascade, the functions of Enzymes E2 (∼40 members) and E3 (> 600 members) have been broadly studied due to that they may regulate only a specific or a single aspect of physiology and general loss of one of E2 or E3 members is insufficient to induce cell death (7, 8). However, due to the limited members of E1 (only 2 members, UBA1 and UBA6, in mouse and human) and lethality induced by loss of one of the two E1 members, their function and action mechanisms in the *in vivo* model have not been well documented (9, 10). Interestingly, *UBA6* encodes a cytoplasm isoform for UBA6 while *UBA1* encodes two isoforms: UBA1a and UBA1b. UBA1a is with nucleic signal and locates in the nuclei while UBA1b is shorter in protein length (without the N-terminal nucleic signal) and locates in the cytoplasm (11). Based on conditional knockout (CKO, i.e *NestinCre*-mediated), role of *Uba6* in mouse has been implicated in neuronal cells (12), and the regulation of polyalanine stretch by UBA6 is suggested in the critical biological process (13). However, the *in vivo* role of *UBA1* in mammals remains largely unknown (14).

Although complete loss of *UBA1* or *UBA6* is hypothesized to induce cell death, somatic mutations in these two genes in human clinic samples offer alternative approaches to understanding their functions in eukaryotic cells. It has been reported that a germ-line mutation in *UBA1* (i.e. p.Met539Ile or p.Ser547Gly) induces X-linked infantile spinal muscular atrophy (XL-SMA), a dysfunction of nervous system (15, 16). Additionally, *UBA1^M41L^*mutation-induced loss of function of UBA1b (p.Met41Leu, only the cytoplasm isoform UBA1b was removed while the nucleic isoform UBA1a is unchanged) was implicated in a newly described autoinflammatory haemato-rheumatoid disease VEXAS syndrome reported by *Beck et al* at December 31^st^ of 2020 in *N Engl J Med* at 2020 (17). The auto-inflammatory disease VEXAS syndrome (Vacuoles, E1 enzyme, X-linked, Autoinflammatory, Somatic) took place mostly in adult male patients (female VEXAS syndrome patients were reported rarely), suggesting it is induced by loss-of-function of *UBA1* (18). Since 2021, more than 200 follow-up retrospective clinical studies describing more than 500 VEXAS syndrome patients further demonstrated that various mutations in *UBA1* were associated with this adult-onset severe disease affecting organs including blood, nose, ear, skin and lung (18–33). Loss of *UBA1b* homologs rather than *UBA1a* in zebrafish by morpholino had been reported to induce systemic inflammation in the same study by *Beck et al* (17), but the mutants were lethal and minimally reminiscent of the VEXAS symptoms in human. Recently, through cutting-edge gene editing strategy, a mouse cell line model and a human HSCs-based model (hosted by chimeric mice) for VEXAS syndrome was reported by *Chiaramida et al* (34) in *Blood Advances* and by *Molteni et al* in a meeting abstract respectively (35). Although the mouse cell line and the “the chimeric mice with gene-edited human HSCs” model are important and complementary for understanding pathology of VEXAS syndrome, a genetic and reliable *in vivo* VEXAS-like animal model maintained at the adult ages is still lacking.

To define the major responsible cell type(s) and recapitulate human VEXAS syndrome in mammals, in this study we aimed to generate murine models by genetically manipulating *Uba1* locus and to further explore the pathology and treatment of the autoinflammatory disease. Although at the moment by the gene-editing approach we failed to obtain alive mouse models with an exact point-mutation in *Uba1* in mouse HSCs as that in VEXAS syndrome patients (i.e. *Uba1^M41L^*), we successfully constructed various conditional knockout (CKO) models (9 lines of CKO mutants induced by various Cre strains plus 3 different bone marrow chimeric approaches were included in the study) to dissect the discrete role of *Uba1* in almost each major hematopoietic cell type including HSCs, lymphoid cells, megakaryocytes and myeloid cells. Our study suggests that loss of *Uba1* in neutrophils is critical for inducing VEXAS-like disorders and offer a translational implication suggesting that the VEXAS symptom is likely attributed to a combinational hematological and immunological consequence of *UBA1* point-mutation over a long-term journey of clonal hematopoiesis. Thus, both aberrant HSCs and neutrophils are equally suggested as key drivers of the autoinflammatory disease. In addition, our study also suggested that treatment with IL-1β/IL-1R1 inhibitors Anakinra or Canakinumab or genetic loss of the myeloid pro-survival regulator *Morrbid* partially mitigated the VEXAS-like symptoms in mice.

## RESULTS

### Conservation of UBA1 in mammals and the Cre/flox-based approach to model VEXAS *syndrome* in mouse

*UBA1* in human and mouse is at the X-chromosome and shares the same protein length and almost the same protein molecular weight (1,058 amino acids; 117.849 kDa vs. 117.808 kDa for the full-length isoform). The identity of their UBA1 protein sequences is as high as 95% and similarity of that as high as 98% (**Figure S1A**). Since the first report of VEXAS syndrome at the year 2020, more than two hundred of retrospective clinical studies have been reported in the past 4 years, including several national-wide large cohort reports, suggesting that VEXAS syndrome is generally ignored but a prevalent entity for aged males (1:4000 for males older than 50) (26). Although VEXAS syndrome is induced by a spontaneous somatic mutation in *UBA1* at HSCs in human, here we report results on Cre/flox-based conditional knockout (CKO) models, by observing the primary mutant mice or by transplantation of the bone marrow cells in the chimeric mice for further studies. The outcome suggests the CKO strategy is technically feasible and easy to obtain. In addition, the mutants could be maintained during adult age since we identified neutrophil loss of *Uba1* (null mutation) induces VEXAS-like symptoms (**Figure 1A**). Due to unavailability of certain *Cre* strains, the present study has not covered the role of *Uba1* specific for erythroid blasts, mast cells, eosinophils and basophils; but other major cell types in hematopoiesis system have been covered and described in detail below. Here we report the results from the 9 different CKO mouse models covering deletion of *Uba1* in HSCs (3 lines in total), pan-lymphoid cells (2 lines in total) and pan-myeloid cells (4 lines in total; among them, 1 line for megakaryocyte).

**Figure 1:**
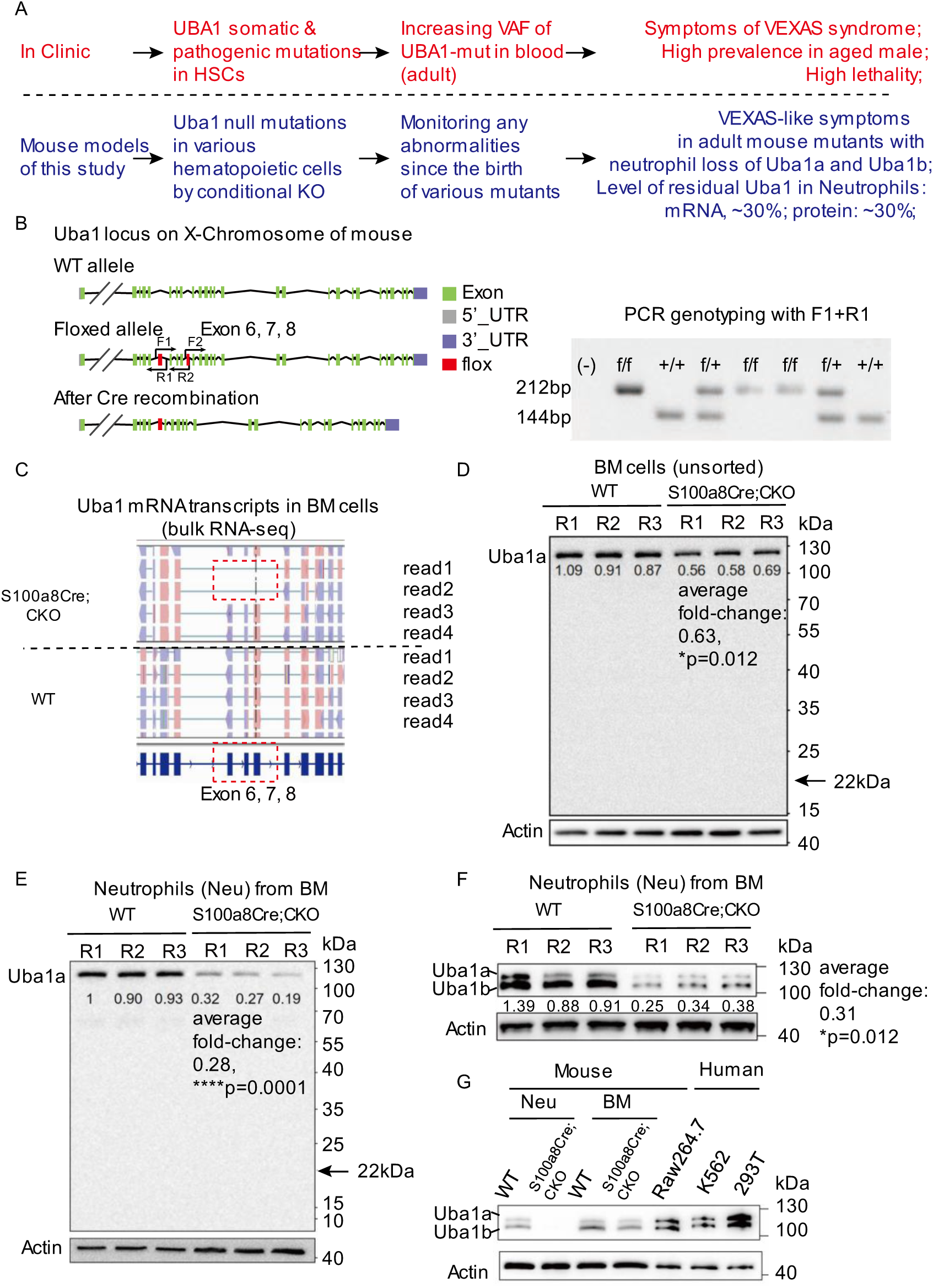
Conditional depletion of the conserved ubiquitin-activation protein Uba1 in mice. **(A)** A brief introduction of VEXAS syndrome occurrence in clinic and in the present study using CKO mouse models. **(B)** Gene structure of *Uba1* in mice and the CKO strategy in the study. Mouse *Uba1* locus is in X-chromosome as human *UBA1*. We generated a flox allele to dissect the role of Uba1. The Exons 6-8 of the gene were depleted by the Cre/flox genetic strategy to generate CKO mutants. Left panel: gene structure of *Uba1* in WT allele and the flox (knockin) allele; Right panel: primers F1/R1 were used to distinguish WT and flox allele in the PCR genotyping. **(C)** Depletion of *Uba1* transcription was revealed by sequencing-read alignment in the bulk RNA-seq datasets (the analysis tool: the IGV software). Note that in the cropped frame, 2 out of 4 sequencing-reads are broken (with depletion of Exons 6-8) in CKO mice while all of sequencing-reads are intact in WT, consistent with the flox design shown in **B**. The CKO mice used here for transcriptional chimerism analysis are *S100a8Cre-CKO*. For the VEXAS-like phenotype of *S100a8Cre-CKO* see Figure 4. **(D)** Expression of Uba1 protein in total BM cells by an antibody specific to Uba1a isoform. Note the residual level of Uba1a in the unsorted BM cells of *S100a8Cre-CKO* is about 63% of that in WT controls (p=0.012). In addition, no truncated Uba1a protein is detected (arrow, predicted molecular weight 22kDa). Three biological repeats were used for WT and CKO mice (R1, R2 and R3). **(E)** Expression of Uba1 isoform in purified neutrophil cells by the same Uba1a-specific antibody. Note the residual level of Uba1a in isolated neutrophils of *S100a8Cre-CKO* is about 28% of that in WT controls (p=0.0001). Again, no truncated Uba1a protein is detected (arrow, predicted molecular weight 22kDa). **(F)** Neutrophil depletion of Uba1a and Uba1b determined by an alternative antibody sensitive to both isoforms. Note the residual level of Uba1 (two isoforms in total) in *S100a8Cre-CKO* is about 31% of that in WT controls (p=0.012). **(G)** Expression of the two Uba1 isoforms (Uba1a and Uba1b) in primary bone marrow (BM) cells of WT and *S100a8Cre-CKO*, in a mouse cell line Raw264.7 and in two human cell lines K562 and 293T. Note that mouse and human both expresses two isoforms of Uba1 protein: Uba1a and Uba1b. In addition, the Uba1b isoform in the primary mouse BM cells appears to have higher expression level detected by the common antibody. However, the two isoforms were with comparable level in the mouse and human cell lines. *, p<0.05; ***, p<0.001; n=3∼6 biological repeats. R1/R2/R3 indicates three independent biological repeats.

As shown in the left panel of **Figure 1B**, we generated a flox knockin strain at the *Uba1* locus where the Exon 6-8 will be removed when the *Uba1^flox/+^* mice were crossed with certain *Cre* lines. The flox line appears normal as WT mice in male and female examined (for male mice: *Uba1^flox/y^*is normal as *Uba1^+/y^*; and for female mice: *Uba1^flox/flox^* and *Uba1^flox/+^* are normal as *Uba1^+/+^*). PCR genotyping with primers F1 and R1 successfully distinguished the flox knockin allele and wildtype (WT) allele (right panel, **Figure 1B**). As the VEXAS syndrome is a disorder mostly affecting aged men, we mainly generated male CKO mutant mice (*Cre; Uba1^flox/y^*) and collected the tissue samples from the male mutant mice for the following studies described below.

As the CKO mice generally manifest chimerism of WT and mutant mRNA transcripts of *Uba1* in certain tissues, we first verified the deletion of the *Uba1* transcripts by aligning the bulk sequencing reads in the pool of all readouts to the mouse reference genome. As shown in the **Figure 1C**, we observed that some transcripts indeed have no Exon 6-8, which is expected by the Cre/flox targeting strategy (the CKO mice used for the transcriptional chimerism analysis are *S100a8Cre-CKO*). Removal of Exon 6-8 by Cre results in 331-bp off and an early occurrence of a stop codon at the Exon 10 in the mRNA transcript. The predicated truncated protein is 200aa in protein length and about 22kDa in protein molecular weight. Based on the Uba1a-specific antibody, we failed to identify a real truncated protein in the BM cell samples or in the neutrophils of the *S100a8Cre-CKO*. Using BM cells for measuring the residual Uba1 level, we determined the average level of Uba1 in the *S100a8Cre-CKO* mutant is 63% of that in WT controls (p=0.012, **Figure 1D**). Using purified neutrophil cells for measuring the residual Uba1 level, we determined the average level of Uba1 in the *S100a8Cre-CKO* mutant is 28% of that in WT controls (p=0.0001, **Figure 1E**). We determined a similar residual level of Uba1 using an alternative antibody sensitive to both isoforms Uba1a and Uba1b (∼31% of WT level, p=0.012, **Figure 1F**). Although this common antibody used in the study appears to be more sensitive to Uba1b isoform in mouse BM cells, Uba1a isoform is detectable but its expression is weaker than that of Uba1b. In the mouse and human cell lines, Uba1a and Uba1b appear to have similar expression level (**Figure 1G**). We also conducted qRT-PCR assays and determined that residual levels of *Uba1* mRNA transcripts are about ∼50% of that in the total BM cells and ∼30% of that in neutrophils (**Figure S1B**). Of note, although there is slight leakage of *S100aCre* in monocytes determined by qRT-PCR (fold change of mRNA: 0.78, p<0.001; **Figure S1B**), the protein level of residual Uba1 appear grossly normal in the *S100a8Cre-CKO* mutant (fold change of protein: 0.87, p=0.1979; **Figure S1C**). Taken together, the results suggest that mouse *Uba1* is highly conserved and our conditional depletion of *Uba1* transcripts and proteins by the Cre/flox approach is successful. Specifically, in the *S100a8Cre-CKO* mutants, neutrophil-depletion of Uba1a and Uba1b were determined by experimental assays on mRNA transcripts and protein isoforms respectively (residual level is about 30% of that in the WT control).

### Depletion of Uba1 in HSCs induces extensive cell death of hematopoietic cells

As shown in **Figure 2A** and **B**, we tried three different HSCs-targeting *Cre* lines to dissect the role of *Uba1* at the top of the hematopoietic system: *Vav1Cre*, *Rosa26-Cre-Ert2* (labeled as R26CreErt2 for simplicity) and *Mx1Cre*. *Vav1Cre* is active as early as embryonic day 9 (36). After several rounds of breeding, we failed to obtain any male *Vav1Cre; Uba1^flox/y^* pups (**Figure 2A**, total male pups n=16, *Vav1Cre; Uba1^flox/y^* male pups n=0; chi-square test, p<0.05). We then turned to crossbreed *R26CreErt2; Uba1^flox/y^*or *Mx1Cre; Uba1^flox/y^* as they are alive when no drug induction is applied. To dissect the role of *Uba1* in hematopoiesis, we constructed several competitive bone marrow transplantation (cBMT) assays as shown in **Figure 2B**. Donor *Uba1^flox/y^* mice were used as control along with donor mice *R26CreErt2; Uba1^flox/y^* or *Mx1Cre; Uba1^flox/y^.* Of note, *R26CreErt2* is expressed in all cell types upon Tamoxifen induction (37) while *Mx1Cre* is widely used for dissecting role of genes in hematopoietic stem cells upon inflammation induction (i.e. polyI:C) (38). As shown in **Figure 2C** to **H**, activation of the *R26CreErt2* by tamoxifen or activation of *Mx1Cre* by PolyI:C induced cell death for a large fraction of *R26CreErt2; Uba1^flox/y^*or *Mx1Cre; Uba1^flox/y^* donors, suggesting that *Uba1* is required for hematopoietic stem cell and progenitor cell (HSPCs), which are at the top hierarchy level of the hematopoietic system. We were also interested in how long the primary *Mx1Cre; Uba1^flox/y^*mice can survive after PolyI:C induction. As shown in **Figure 2I**, after administration of PolyI:C, the animals cannot survive longer than 3 weeks (tested animals n=5 per group, p<0.001). As *Mx1Cre* is also active in erythroid progenitor cells and the half-life of red blood cells is very short (∼24 hours), one of the direct reasons for the death of *Mx1Cre; Uba1^flox/y^* mutants is probably the shortness of red cell supply (an Erythroid line of *Uba1* depletion is demanded to validate this hypothesis). In conclusion, these results demonstrated that a null version of *Uba1* at the HSC level induces a quick and extensive death in almost all kinds of hematopoietic cells.

**Figure 2:**
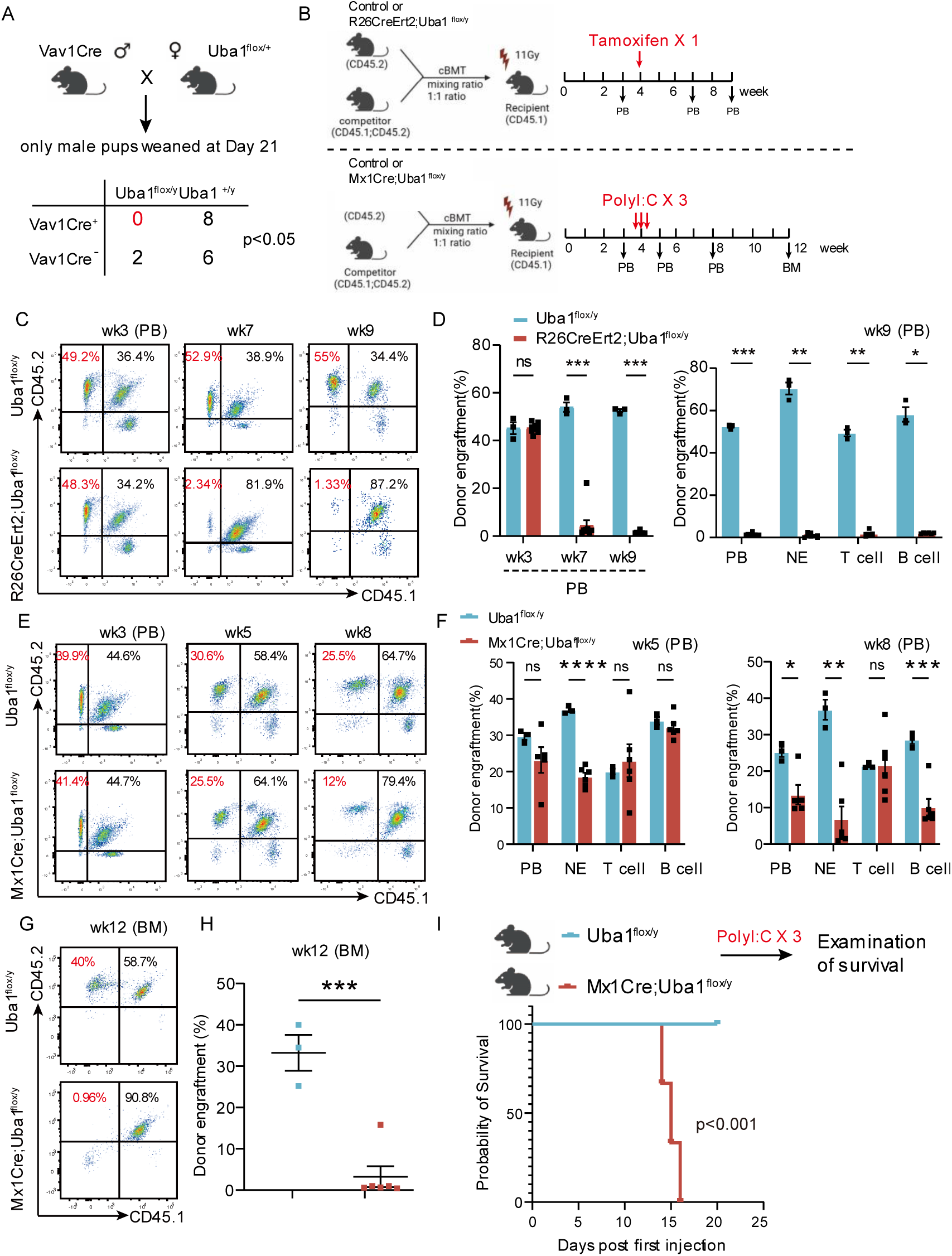
Conditional depletion of *Uba1* in hematopoietic stem cells (HSCs) failed to induce VEXAS-like symptoms in chimeric mice but resulted in a quick and extensive death of a large portion of mutant hematopoietic cells. **(A)** Genetic crossing strategies for generating CKO mutants with depletion of *Uba1* in HSC by *Vav1Cre*, *R26CreErt2* and *Mx1Cre* respectively. In the scheme, the strategy for generation *Vav1Cre; Uba1^flox/y^* is shown. *Vav1Cre* is expressed in HSC as early as embryonic day 10. After collecting male pups from several round of animal birth (n=16 pups), we never obtained the compound mutants *Vav1Cre; Uba1^flox/y^* (p=0.0186, chi-square test), suggesting the *Vav1Cre; Uba1^flox/y^* embryos are lethal at the embryonic stages. **(B)** Inducible depletion of *Uba1* in chimeric mice with BM reconstituted by competitive bone marrow transplant (cBMT) through mixing *R26CreErt2; Uba1^flox/y^* or *Mx1Cre; Uba1^flox/y^* donors (CD45.2^+^) with competitor donors (F1 mice: CD45.1^+^CD45.2^+^) as indicated (mixed ratio, 1:1) while the recipient mice were CD45.1^+^. *Uba1^flox/+^* female mice were crossed with *R26CreErt2* and *Mx1Cre* males to generate the donor mice *R26CreErt2;Uba1^flox/y^*or *Mx1Cre;Uba1^flox/y^*. Induced depletion of *Uba1* in *R26CreErt2;Uba1^flox/y^* chimeric mice was conducted by feeding with Tamoxifen (one dose at the 4th week post cBMT) while induced depletion of *Uba1* in *Mx1Cre; Uba1^flox/y^* chimeric mice was conducted by i.p. injection of PolyI:C (three doses at the 4th week post cBMT). Confirmation of the BM reconstitution is performed using flow cytometry of PB at the third week post cBMT (See **C**). (**C-D**) Representative flow cytometry profiles and the quantification of the donor engraftment in the *R26CreErt2;Uba1^flox/y^* chimeric mice at the indicated time points. N=3∼5 chimeric animals per group. Note that CD45.2^+^ percentage was normal at week-3 (wk3) but dramatically dropped at week-7 (wk7) and week-9 (wk9) post the cBMT. The cBMT experiments with the donors *R26CreErt2; Uba1^flox/y^* were repeated three times and the representative one was shown here. (**E-H**) Representative flow cytometry profiles and the quantification of the donor engraftment in the *Mx1Cre;Uba1^flox/y^* chimeric mice at the indicated time points. **E** and **F** are results from flow cytometry analysis on peripheral blood cells (PB) while **G** and **H** are results from flow cytometry analysis on BM cells. N=3∼6 animals per group. The cBMT experiments with the donors *Mx1Cre;Uba1^flox/y^* were repeated twice and the representative one was shown here. (**I**) *Mx1Cre;Uba1^flox/y^* primary mice proceeded to a quick death within 20 days after induction with PolyI:C. The survival test for *Mx1Cre;Uba1^flox/y^*primary mice was performed only once. Age of the mice, 8∼12 weeks old. N=5 animals per group. ns, not significant; *, p<0.05; **, p<0.01; ***, p<0.001; ****, p<0.0001; n=3∼6 biological repeats.

### Depletion of Uba1 in lymphoid cells or megakaryocytes results in cell death of lymphoid cells or less platelets respectively

We then tested depletion of *Uba1* at the mature blood cells in the hematopoietic and immune system. Two *Cre* lines for lymphoid cells (*Cd4Cre* and *Cd19Cre*) (39, 40), and one *Cre* line for megakaryocytes (*Pf4Cre*) line (41) were used. As shown in **Figure 3A** to **D**, dramatically decreased amount of Cd4^+^T cells or Cd19^+^B cells were readily observed in *Cd4Cre; Uba1^flox/y^* and *Cd19Cre; Uba1^flox/y^* mice (50%∼70% depletion efficiency). However, the mutant animals appear normal and survive over than 12 months in our specific-pathogen-free (SPF) facility. As shown in **Figure 3E** and **F**, ∼50% depletion of megakaryocytes and platelets were also observed in *Pf4Cre; Uba1^flox/y^* but the mutant animals appear normal and no VEXAS-like symptoms were observed. These results demonstrated that depletion of *Uba1* in lymphoid or megakaryocytes failed to induce VEXAS-like abnormalities in mice.

**Figure 3:**
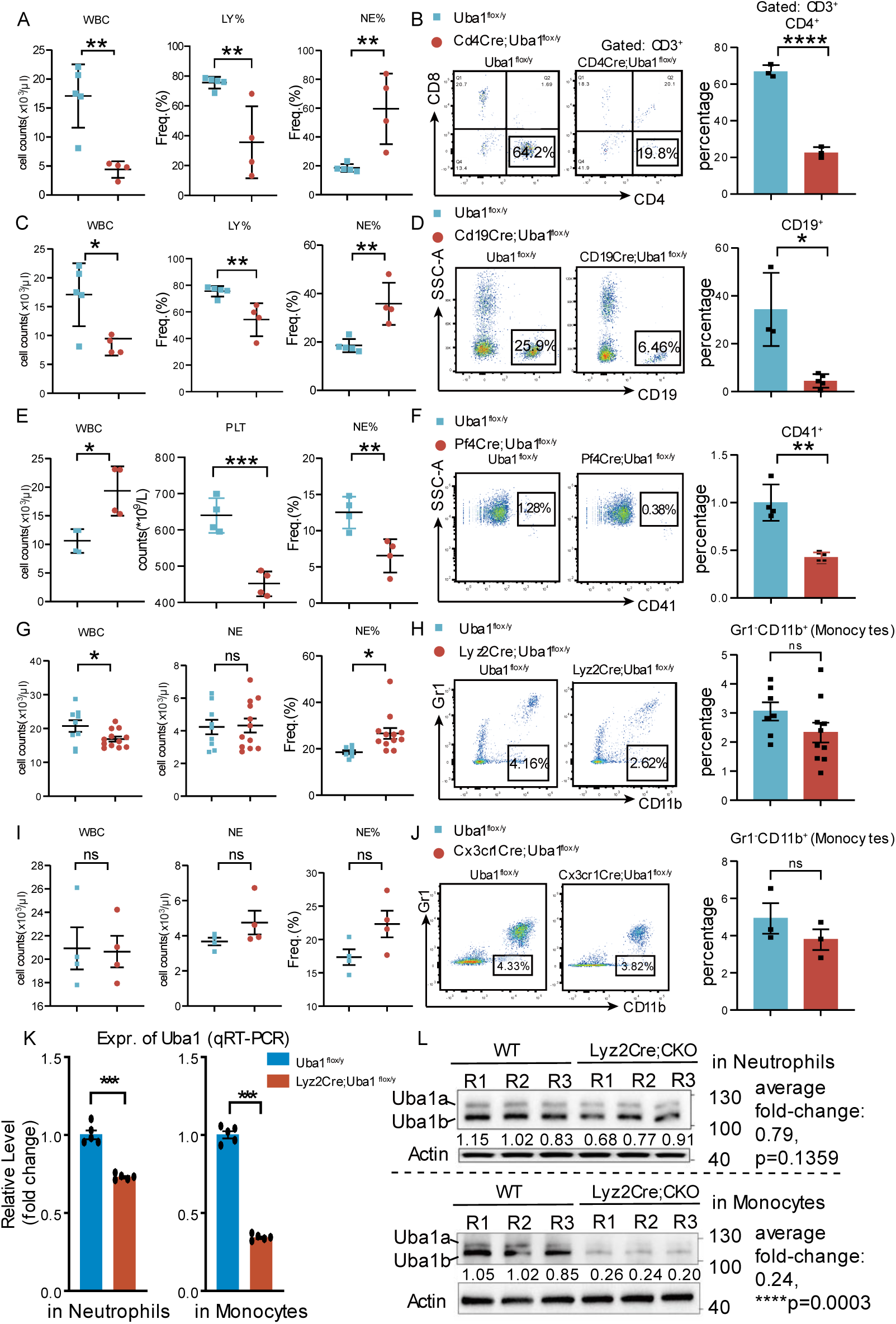
Conditional depletion of *Uba1* in mature CD4^+^T cells, CD19^+^B cells, megakaryocytes and monocytes all failed to induce VEXAS-like symptoms. *Uba1^flox/+^* female mice were crossed with these *Cre-*line male mice to generate the target male mice as indicated in (**A-F**). (**A-D**) Loss of *Uba1* in lymphoid cells (*Cd4Cre*- or *Cd19Cre*-induced) results in obviously decreased number of leukocytes in PB, which were verified by flow cytometry respectively, while the mutant animals were viable and failed to manifest VEXAS-like symptoms. **A** and **B**, results from *Cd4Cre; Uba1^flox/y^* primary male mice; **C** and **D**, results from *Cd19Cre; Uba1^flox/y^* primary male mice. (**E-F**) Loss of *Uba1* in megakaryocytes (*Pf4Cre*-induced) results in decreased number of platelets and megakaryocytes in PB. We verified the decreased megakaryocytes by flow cytometry analysis on PB samples (Cd41^+^). The mutant CKO animals are viable and failed to manifest VEXAS-like symptoms. (**G-J**) Loss of *Uba1* in monocytes/macrophages were viable and failed to induce VEXAS-like symptoms. For dissecting the role of *Uba1* in monocytes, macrophages and microglia cells, we tried two *Cre* lines: *Lyz2Cre* and *Cx3cr1Cre*. These two lines were well characterized and defined in previous studies for the homeostasis in monocytes, macrophages, or microglia cells in the laboratory tests. **G-H**, results from *Lyz2Cre;Uba1^flox/y^*primary male mice; **I-J**, results from *Cx3cr1Cre; Uba1^flox/y^*primary male mice. *Lyz2Cre;Uba1^flox/y^* CKO mutants manifest a subtle increased count of WBC and increased percentage of neutrophils (**I**), detectable but rare vacuoles in myeloid cells from PB. However, the *Lyz2Cre;Uba1^flox/y^* CKO mutants appear quite normal. Similarly, *Cx3cr1Cre; Uba1^flox/y^* CKO mutants also appear quite normal. (**K**) Quantification of residual *Uba1* mRNA transcripts by qRT-PCR assays. Residual average level of *Uba1* mRNA transcripts in purified neutrophils mutants is about 75% while that in the purified monocytes is 25% in *Lyz2Cre;Uba1^flox/y^* CKO compared to the controls. N=5 biological repeats. (**L**) Quantification of residual Uba1 proteins by western blotting assays. Upper panel, expression of Uba1 in isolated neutrophils; Lower panel, expression of Uba1 in isolated monocytes. Note that in the *Lyz2Cre;Uba1^flox/y^*CKO mutants, greater than 70% reduction of Uba1 in monocytes (average residual level: 24%, p = 0.0003) while less than 30% reduction of Uba1 (average residual level: 79%, p = 0.1359) were detected at the protein level. R1/R2/R3 indicates three independent biological repeats. ns, not significant; *, p<0.05; **, p<0.01; ***, p<0.001; ****, p<0.0001; n=3∼11 biological repeats.

### Mice with depletion of Uba1 in monocytes by Lyz2Cre *or* Cx3cr1Cre appear normal

We also tested depletion of *Uba1* in monocytes and macrophages. For this aim, two Cre lines for monocytes or macrophages were used in the study: *Lyz2Cre* and *Cx3cr1Cre* line (42, 43). In contrast to *Pf4Cre; Uba1^flox/y^, Cd4Cre; Uba1^flox/y^* and *Cd19Cre; Uba1^flox/y^*, as shown in **Figure 3G** to **J**, no deduction of myeloid cells was observed in *Lyz2Cre; Uba1^flox/y^* and *Cx3cr1Cre; Uba1^flox/y^*. Although we occasionally observed vacuoles in few myeloid cells from the blood cells of *Lyz2Cre; Uba1^flox/y^*, both *Lyz2Cre; Uba1^flox/y^* and *Cx3cr1Cre* mice appear quite normal. To confirm the *Cre* activity induced by *Lyz2Cre*, we crossed *Lyz2Cre; Uba1^flox/y^* with *tomato-lox-GFP* line (labeled as *lox-GFP* here for simplicity) (44) for generating *Lyz2Cre; lox-GFP* and *Lyz2Cre; lox-GFP; Uba1^flox/y^*. Expression of GFP in *Lyz2Cre; lox-GFP* and *Lyz2Cre; lox-GFP; Uba1^flox/y^* indicates the activity of *Lyz2Cre* and the depletion of *Uba1* in the *Lyz2Cre; lox-GFP; Uba1^flox/y^* compound line. As shown in **Figure S1D** and **E**, surprisingly loss of *Uba1* even resulted in a higher portion of myeloid cells in the peripheral blood. To further characterize the myeloid cell-depletion of *Uba1* by *Lyz2Cre*, we performed the qRT-PCR and immunological blotting assays. As shown in **Figure 3K** and **L**, a significant deduction of Uba1 is detected in the monocytes of *Lyz2Cre; Uba1^flox/y^*by both qRT-PCR and immunological blotting assays (residual level is about 25%, p<0.001). A slight deduction of Uba1 is also detected in neutrophils of *Lyz2Cre; Uba1^flox/y^*compared to the controls (residual level is about 75% by qRT-PCR, p<0.001 and about 78%, p=0.116), suggesting slight leakiness by Lyz2Cre in neutrophils in the mutants. Taken together, these results demonstrated that monocyte-based depletion of *Uba1* by *Lyz2Cre* or *Cx3cr1Cre* failed to induce VEXAS-like abnormalities in mice, and intriguingly somehow myeloid cells with loss of *Uba1* appear to have extended life-span.

### Depletion of Uba1 in neutrophils induces VEXAS-like phenotypes

Although *Lyz2Cre* demonstrate slight leakiness in neutrophils, it is not a strong and reliable *Cre* stain for neutrophils. We then tried depletion of *Uba1* in neutrophils using the *S100a8Cre* line (56). Most of the *S100a8Cre; Uba1^flox/y^* mutant mice (for simplicity, we sometime label the mice as *S100a8Cre-CKO* in the study) appear normal at the age 2∼3-month-old while a few *S100a8Cre-CKO* mice appear abnormal as early as 1-month old (frequency: 5 out of 47). When the *S100a8Cre-CKO* mutant mice grew up (about 4-month-old) in the specific-pathogen-free (SPF) facility, we started to observe their kink tails (**Figure 4A**) and flare nose (**Figure 4B**). And such aberrations turned to be more obvious at older ages (i.e 6∼12-month-old). Additionally, hair loss on their back started to be observed (**Figure 4A**), a sign of dermatitis. Furthermore, after examining the entire body, we also observed swollen toes or pigmentation on the toes of the mutant mice (**Figure 4C**). At the moment we don’t know the exact mechanism(s) by which the pigmentation appears in the mutant toes and fingers.

**Figure 4:**
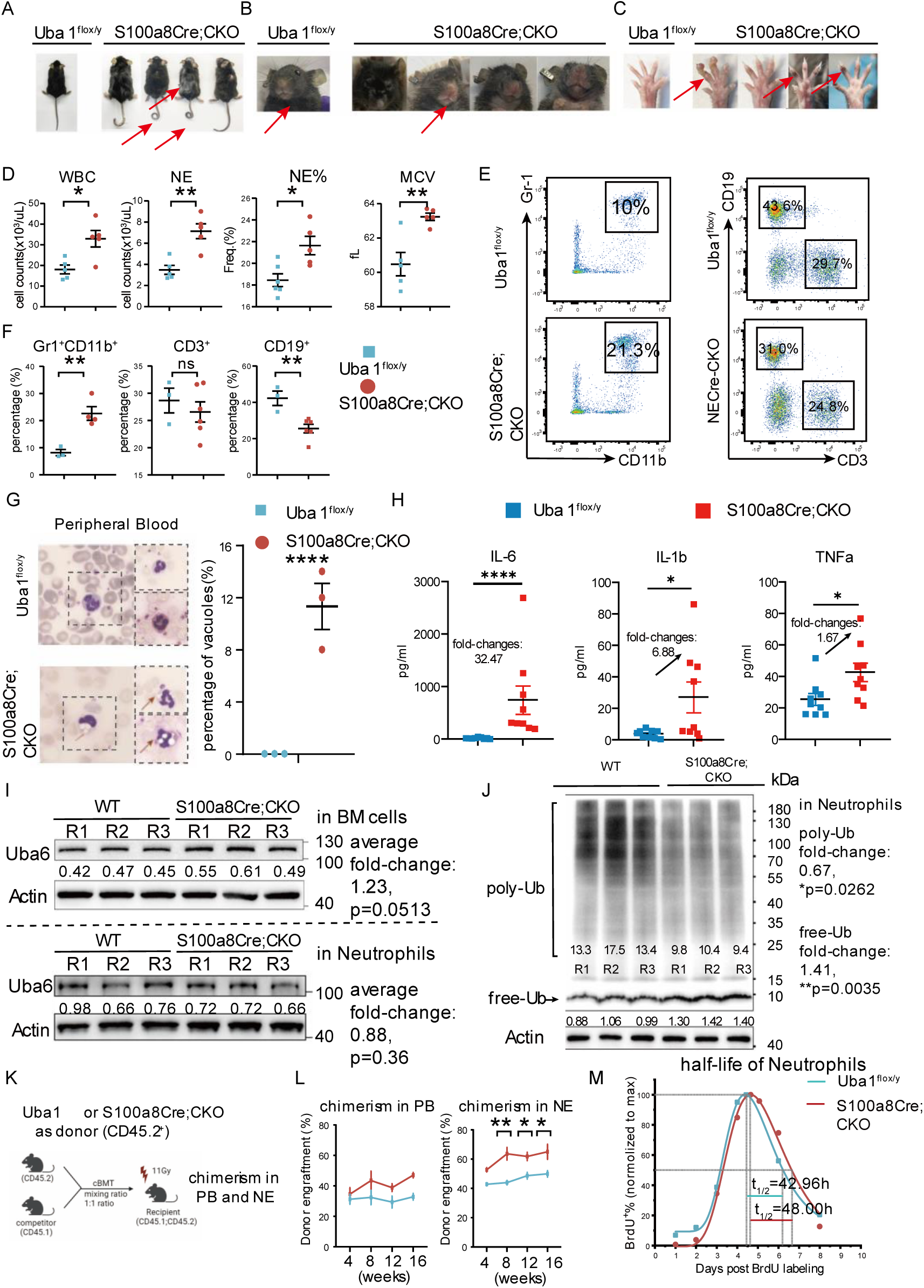
Conditional depletion of *Uba1* in neutrophil cells induce VEXAS-like auto-inflammatory symptoms. Similarly, *Uba1^flox/+^* female mice were crossed with *S100a8Cre* male mice to generate the target male mutant mice *S100a8Cre;Uba1^flox/y^* (*S100a8Cre;CKO*). Of note, female *S100a8Cre;CKO* were also generated in the study and had similar phenotypes as the male *S100a8Cre;CKO* described below (n=10 for female mice; n>40 for male mice). For a study comparable to human VEXAS Syndrome, we only report the data from male *S100a8Cre;CKO* mutants. (**A**) *S100a8Cre;CKO* mice manifest obvious dermatitis and hair loss on their back (penetration rate: 100%; n=20). (**B**) *S100a8Cre;CKO* mice manifest flare noses (penetration rate: 100%; n=20). (**C**) *S100a8Cre;CKO* mice manifest swollen paw (penetration rate: ∼50%; n=20) with pigmentation on the skin of their fingers (penetration rate: 100%; n=20). (**D**) Aberrant hematological parameters in PB of *S100a8Cre;CKO* mutants. Consistent to couple of the most sensitive laboratory tests for VEXAS, increased counts of white blood cells (WBC) and neutrophils were observed; Of note, a significant increase of mean corpuscular volume (MCV) is also determined in *S100a8Cre;CKO* mutants. Penetration of aberrant hematologic parameters is age-dependent. At older age (6∼12-month-old), the penetrance is about 100%, at the younger age (3∼5-month-old), the penetrance is about 60%. Mice used here are at older age. (**E-F**) Representative flow cytometry profiles of PB and quantifications of mature neutrophils (Gr1^+^CD11b^+^) and lymphoid cells (CD3^+^ or CD19^+^). (**G**) Vacuolar neutrophils were identified in the *S100a8Cre;CKO* mice by Giemsa staining but not in the WT controls. The frequency is not as high as in human VEXAS patients as the rate we quantified is only ∼10% in *S100a8Cre;CKO* mice but undetectable (0%) in WT controls. Fifty cells with nuclei on the Giemsa slides (n=3 biological repeats) were randomly chose to identify if vacuoles exist or not. (**H**) Serum level of pro-inflammatory cytokines IL-6, IL-1β, and TNFα all were upregulated in the VEXAS-like mutant *S100a8Cre;CKO* mice. (**I**) Expression of another E1 enzyme Uba6 appears grossly normal in total BM cells (upper panel) and isolated neutrophils (lower panel) of the *S100a8Cre;CKO* mice compared to that of the control mice. R1/R2/R3 indicates three independent biological repeats. (**J**) Poly-ubiquitin (poly-Ub) and free ubiquitin (free-Ub) levels were quantified by western blotting in isolated neutrophils of the WT and *S100a8Cre-CKO*. We observed significant reduction of poly-Ub (fold change: 0.67, p=0.0262) and a significantly increased level of free-Ub (fold change: 1.41, p=0.0035) in the purified neutrophils of *S100a8Cre;CKO* mice compared to that of WT controls. R1/R2/R3 indicates three independent biological repeats. (**K-L**) Comparing the growth or survival advantage of WT- and mutant-neutrophils by cBMT analysis of bone marrow (BM) cells from *S100a8Cre;CKO* mice and WT donors. **K**, the scheme for the experimental procedure using cBMT assays. **L**, chimerism analysis in PB or in neutrophils at the indicated time points post the cBMT. (**M**) Measuring the half-life of neutrophils in the peripheral blood by BrdU pulse assays. See Methods for details about the BrdU staining, flow cytometry and computational analysis. ns, not significant; **, p<0.01; n=3∼20 biological repeats.

When we extended the production of *S100a8Cre; Uba1^flox/y^*male mutant mice by genetic breeding, we were able to obtain *S100a8Cre; Uba1^flox/flox^* female mutant mice in the offspring. The female mutant mice manifest identical appearances as that described above in the male mice, and the controls including the *S100a8Cre* line itself (male or female) or *Uba1^flox/y^* male or *Uba1^flox/flox^*female are as normal as wild type (WT) when aged. The male and female mutant mice are fertile but the overall fertility appeared to be disturbed to certain extent (the disturbed fertility has not yet been quantified yet and will be examined in more rigorous control settings in future studies). In addition, we alternatively husbanded the mutant male or female mutant mice in a clean-grade air condition rather than the SPF-grade facility. The mutant mice in the clean-grade facility showed similar symptoms as that in the SPF facility and no obvious pneumonia or colitis were observed (N=5). Taken together, these phenotypic observations confirmed that the appearances of kink tails, flare noses, and pigmentations on paws were induced by *S100a8Cre*-induced depletion of *Uba1* in the mutant male and female mice and that husbandry of the mutant mice in the SPF-grade or clean-grade facility manifest similar symptoms mentioned above.

To quantify any aberrant hematological parameters in the mutant mice with similarities to the VEXAS syndrome in human, we quantified the blood cells by haemato-meters and flow cytometry. As shown in **Figure 4D**, the *S100a8Cre; Uba1^flox/y^*male mice manifest increased white blood cell (WBC) counts (p<0.05), increased neutrophil (NE) counts (p<0.01) and increased neutrophil percentage (NE%, p<0.05), which are hallmarks and indications of autoinflammatory diseases including VEXAS syndrome. Furthermore, a significant increase of mean corpuscular volume (MCV) was also determined in *S100a8Cre-CKO* mutants (**Figure 4D**). Of note, MCV is recognized as one of the most sensitive laboratory tests for VEXAS syndrome and increase of MCV indicates anemia in clinic (17). The differences in red blood cell (RBC) counts between *S100a8Cre-CKO* mutants and the WT controls did not reach the bar of statistical significance in the *S100a8Cre-CKO* mutants compared to the controls. We also used flow cytometry to measure the changes in compartments of mature blood cells the *S100a8Cre-CKO* mutant mice or hematopoietic progenitor cells in bone marrow (BM). As shown in **Figure 4E-F**, neutrophil’s fraction (Gr1^+^CD11b^+^) is increased in PB while lymphoid fraction (B cells, CD19^+^) is decreased, consistent to results from haemato-meters (**Figure 4D**).

Compared to WT controls, fraction of HSCs in the mutants is comparable while that of granulocyte-macrophage-progenitor (GMP) and Lin^−^Sca1^−^cKit^+^ cells (LSK) appears to be elevated (**Figure S2A** and **B**). We also examined the existence of vacuoles in the myeloid cells in the PB of the *S100a8Cre-CKO* mutants, one of the hallmarks of VEXAS syndrome. As shown in **Figure 4G**, we observed ∼10% of myeloid cells (mainly are neutrophils) are with vacuolar characteristics. H&E staining of skin, tail and paws indicate dermatitis and infiltration of inflammatory immune cells in the paws (**Figure S3A-C**). However, we did not observe chondritis in the nose in the *S100a8Cre-CKO* mice (relapsing polychondritis in nose and ear is another one of hallmarks of VEXAS syndrome in human). H&E staining of lung tissue identified minimal alteration of immune cells and the stromal cells but single-cell RNA-sequencing profiling detected increased inflammatory scores in the immune cells (**Figure S4**). Single-cell analysis of skin tissue, however, failed to identify obvious alterations probably due to insufficient capture of immune cells in the mutant skins (**Figure S5**). Taken together, we summarize the phenotypic results as that *S100a8Cre-CKO* manifest most of the VEXAS symptoms, including: (1) inflammatory hematological parameters, vacuoles in neutrophils (∼10%), dermatitis, swollen paws; (2) the mutant mice manifest a frequent finger pigmentation which were not reported in human VEXAS syndrome; (3) the mutant mice did not show obvious polychondritis in ear or nose.

We measured the serological parameters in the mutant mice as proinflammatory cytokine level is a key for the autoinflammatory disease VEXAS syndrome. A 10-cytokine panel was used for quantifying proinflammatory and anti-inflammatory cytokines. Among them, three proinflammatory cytokines IL-1β, IL6 and TNFα all are significantly upregulated in the serum with fold change between 2 and 40 accordingly (**Figure 4H**, n=9; see **Figure S6A** for the full plots of the 10 cytokines in the tested mutant animals). Interestingly we observed positive correlation between the three proinflammatory cytokines (IL1β, IL6 and TNFα) but IFNγ appear unchanged (**Figure S6B**), suggesting the three classical proinflammatory cytokines are systematically coordinated in the VEXAS-like mice. These results demonstrate that the mutant mice manifest obvious autoinflammatory haemato-rheumatoid disorders serologically.

We also confirmed grossly normal expression of Uba6 in total BM cells and neutrophils but significantly down-regulated poly-Ubiquitylation (poly-Ub) level and up-regulated free-Ubiquitin (free-Ub) in the mutant neutrophils from the *S100a8Cre-CKO* mice (**Figure 4I** and **J**). Furthermore, the increased count of neutrophils in the *S100a8Cre-CKO* mutants is intriguing as the aberration is in line with what we observed in the *Lyz2Cre; Uba1^flox/y^* mice. Profiling the hematopoiesis by flow cytometry also suggested a biased myelopoiesis (myeloid-skewing) in the *S100a8Cre-CKO* mice (**Figure S2A-B**). In addition, the body weight of the *S100a8Cre-CKO* mutant mice is less than that of WT controls while the weight of spleen is greater than that of WT controls suggesting a splenomegaly in the mutant mice (**Figure S2C**).

Although in neutrophil we observed a slightly increased apoptosis activity determined by Annexin-V-marked flow cytometry (**Figure S2D**), we demonstrated a survival advantage of *S100a8Cre-CKO* myeloid cells (especially neutrophils) by three independent approaches. <u>The first approach</u> is similar to what shown in *LyzCre;lox-GFP* (**Figure S1D** and **E)**, using the GFP reporter line along with flow cytometry analysis. The GFP reporter experiments suggest that a comparable or even higher fraction of Gr1^+^ cells or GFP^+^ cells in the *S100a8Cre-CKO;lox-GFP* compared to that in the *S100a8Cre;lox-GFP* control (**Figure S2E** and **F**). <u>The second approach</u> is through competitive bone marrow transplantation (cBMT) assays (**Figure 4K**). As shown in **Figure 4L**, the donors from *S100a8Cre-CKO* manifest a phenotype of clonal hematopoiesis as shown in the human (45), suggesting that the *S100a8Cre-CKO* neutrophils indeed have a survival advantage somehow. The cBMT chimeric mice (50% *S100a8Cre; Uba1^flox/y^* donor plus 50% *Uba1^flox/y^* donor) manifest subtle VEXAS-like symptoms at the time point 5 months post the cBMT. However, the chimeric mice with 100% *S100a8Cre-CKO* donor manifest similar symptoms as the primary *S100a8Cre-CKO*, suggesting the VEXAS-like symptoms in *S100a8Cre-CKO* were attributed to aberrations in hematopoietic cells and were transplantable when the mutant bone marrow cells were repopulated. <u>The third approach</u> is to directly measure the half-life of neutrophils by BrdU staining and tracing (46). As shown in **Figure 4M**, the mutant neutrophils manifest slightly extended half-life compared to the wild type neutrophils (*t_1/2_* in mutant neutrophils = 48.00 hrs vs. *t_1/2_* in WT neutrophils = 42.96 hrs). Taken together, using three independent *in vivo* experimental assays, we demonstrated that the mutant neutrophils lacking expression of *Uba1* in the *S100a8Cre-CKO* somehow have growth or survival advantage against the WT controls in the bone marrow system.

### Verification of neutrophil-specific loss of Uba1 by scRNA-seq analysis

As shown in **Figure 1D-F**, we have determined neutrophil depletion of Uba1a and Uba1b at the mRNA and protein level (residual level: ∼30%, indicating 70% depletion efficiency). To unbiasedly and orthogonally verify the exact cell type(s) with loss of *Uba1* expression in *S100a8Cre-CKO*, we took advantage of single-cell RNA-sequencing (scRNA-seq) technologies as it in a high-throughput and unbiased way measures gene expression level at the transcriptomic wise in each cell. As shown in **Figure 5A-C**, a total of 11,204 and 10,054 BM cells were included in the UMAP plot composing cells from 3 *Uba1^flox/y^* donors and 3 *S100a8Cre; Uba1^flox/y^* donors. The major BM cell types were annotated (11 cell types in total, left panel of **Figure 5A**). The fractions of HSPCs (mainly are GMPs) and neutrophils are marked by expression of *Ms4a3* and *Cebpd* (right panel of **Figure 5A**). A large deduction of *Uba1* in Pro_Neutrophils and Neutrophils are detected (residual level: 27% and 29% respectively, indicating ∼70% depletion efficiency in both; **Figure 5B**) along with subtle deduction of *Uba1* in monocytes (residual level: 83%, indicating ∼20% depletion efficiency; **Figure 5B**). We did not detect dramatic deduction of *Uba1* in other cell types including HSPCs (**Figure 5B**). These results provide direct evidence suggesting a profound depletion of *Uba1* in neutrophils of *S100a8Cre; Uba1^flox/y^*.

**Figure 5:**
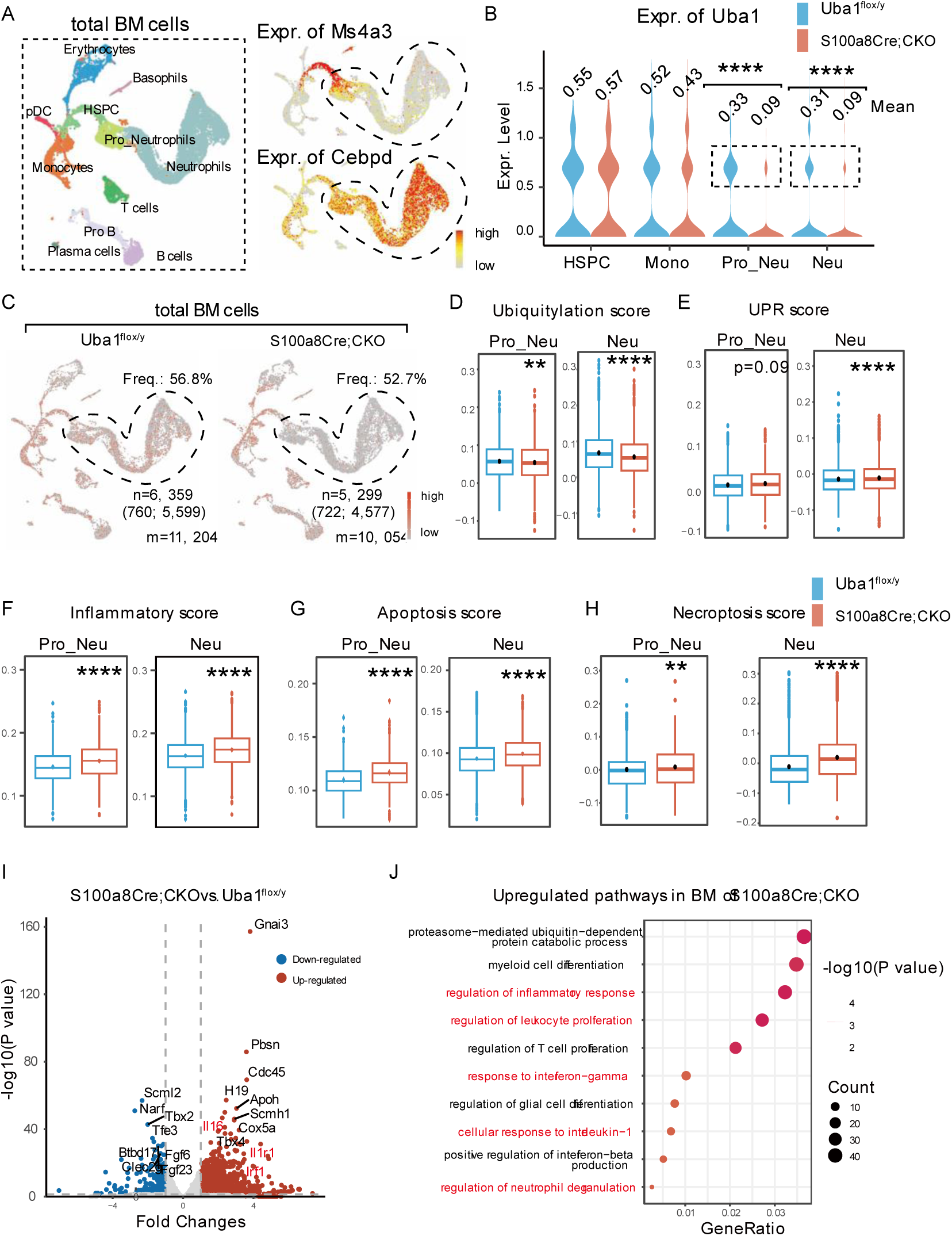
Verification of neutrophil-specific loss of *Uba1* in *S100a8Cre;CKO* by scRNA-seq analysis; and a disturbed neutrophil homeostasis is determined by computational analysis of the single-cell dataset from *S100a8Cre;CKO*. (**A**) BM cells from WT and *S100a8Cre;CKO* mice were subject for single-cell RNA-sequencing (scRNA-seq) analysis to further confirm the neutrophil loss of *Uba1*. Fresh mono-nucleic BM cells (unsorted) were prepared for scRNA-seq. A standard Seurat-based pipeline was conducted to analyze the WT and *S100a8Cre;CKO* scRNA-seq datasets. As shown in the left panel, a total of 11 major hematopoietic cell types were annotated. Of note, two types of neutrophils are highlighted: Pro-Neutrophils and Neutrophils. As shown in the right panel, expression of *Ms4a3* (marker for granulocyte-macrophage progenitor [GMP] cells) and *Cebpd* (marker for primitive and mature neutrophils) marks the developmental continuity from HSPC/GMP to neutrophils. (**B**) Expression of *Uba1* in HSPCs, Monocytes, Pro-Neutrophils and Neutrophils are quantified in WT and *S100a8Cre;CKO* scRNA-seq datasets. Note that there is no deduction of *Uba1* expression in HSPCs from the *S100a8Cre;CKO* (residual level: 104%); a slight decrease of *Uba1* in Monocytes (residual level: 82%, indicating ∼18% depletion efficiency) from the *S100a8Cre;CKO* was determined; a dramatic decrease of *Uba1* in Pro-Neutrophils or in Neutrophils from the *S100a8Cre-CKO* were determined (residual level: 27% and 29%, indicating ∼73% and ∼71% depletion efficiency, respectively). (**C**) Pro-Neutrophils and Neutrophils were highlighted in scRNA-seq datasets and for down-stream analysis of cell homeostasis. For WT, 760 Pro-Neutrophils and 5,599 Neutrophils included in the downstream analysis; For *S100a8Cre;CKO*, 722 Pro-Neutrophils and 4,577 Neutrophils included in the downstream analysis. (**D**) *S100a8Cre-CKO* mice manifest slightly decreased ubiquitination score in Pro-Neutrophils or in Neutrophils compared to WT. The result is expected as Uba1 is an E1 enzyme responsible for ubiquitin activation. (**E-H**) Loss of *Uba1* in neutrophils results in disturbed cellular homeostasis as indicated by increased UPR Score, Inflammatory Score, Apoptosis Score and Necroptosis Score. (**I-J**) In parallel with the scRNA-seq analysis, bulk RNA-sequencing analysis was conducted using total BM cells from WT and *S100a8Cre-CKO*. **I**, volcano plot suggesting some pro-inflammatory genes are upregulated in *S100a8Cre-CKO* mice. **J**, enriched pathways in *S100a8Cre;CKO* mice including pro-inflammation response and cell responses to IL-1 and IFN-gamma. ns, not significant; *, p<0.05; **, p<0.01; ***, p<0.001; ****, p<0.0001. all Pro-Neutrophils (n=722-760) and Neutrophil cells (n=4,577-5,599) were included for the scoring analysis in **D** to **H**. n=3 biological repeats for bulk RNA-seq.

### Disturbed bone marrow cellular homeostasis in the VEXAS-like mutants, revealed by transcriptomic and proteomic analysis

We also took advantage of the scRNA-seq datasets to measure the cell homeostasis by scoring activity of various aspects in the neutrophils using a module-score computational function in the Seurat analysis pipeline (see Methods for details). Cells (Pro_neutrophils and neutrophils) were for scoring analysis are highlighted in **Figure 5C**. As shown in **Figure 5D**, the ubiquitylation scores were readily decreased in Pro_Neutrophils and Neutrophils of the mutants. Scoring the activity of unfolded protein response (UPR), inflammation, apoptosis, necroptosis, reactive oxygen species (ROS), and neutrophil extracellular trap formation further suggested disturbed cellular homeostasis (**Figure 5E-H**). We also cross-validated the results of scRNA-seq using the regular bulk RNA-seq datasets. As shown in **Figure 5I-J**, we detected upregulation of inflammatory genes *Il1r1* and other genes in the inflammatory pathways IL-1 pathway and IFNγ pathway.

In addition, as shown in **Figure S7A-H**, LC/MS-based proteomic analysis of the BM cells from 3 *Uba1^flox/y^* donors and 3 *S100a8Cre; Uba1^flox/y^* donors also demonstrate depletion of Uba1 (This is an orthogonal assay: ∼50% depletion efficiency in total BM cells, suggesting ∼70% depletion efficiency in the neutrophils) and disturbed immune responses including complementary response, inflammatory response and neutrophil extracellular traps (NETs) formation. We examined the phagocytosis capability of the mutant neutrophils by co-culture of neutrophils and GFP-labeled *E. coli* bacteria (See **Figure 6A** for the experimental scheme). As shown in **Figure 6B**, the mutant neutrophils appear to have greater phagocytosis capabilities (n = 3 biological repeats, p<0.01).

**Figure 6:**
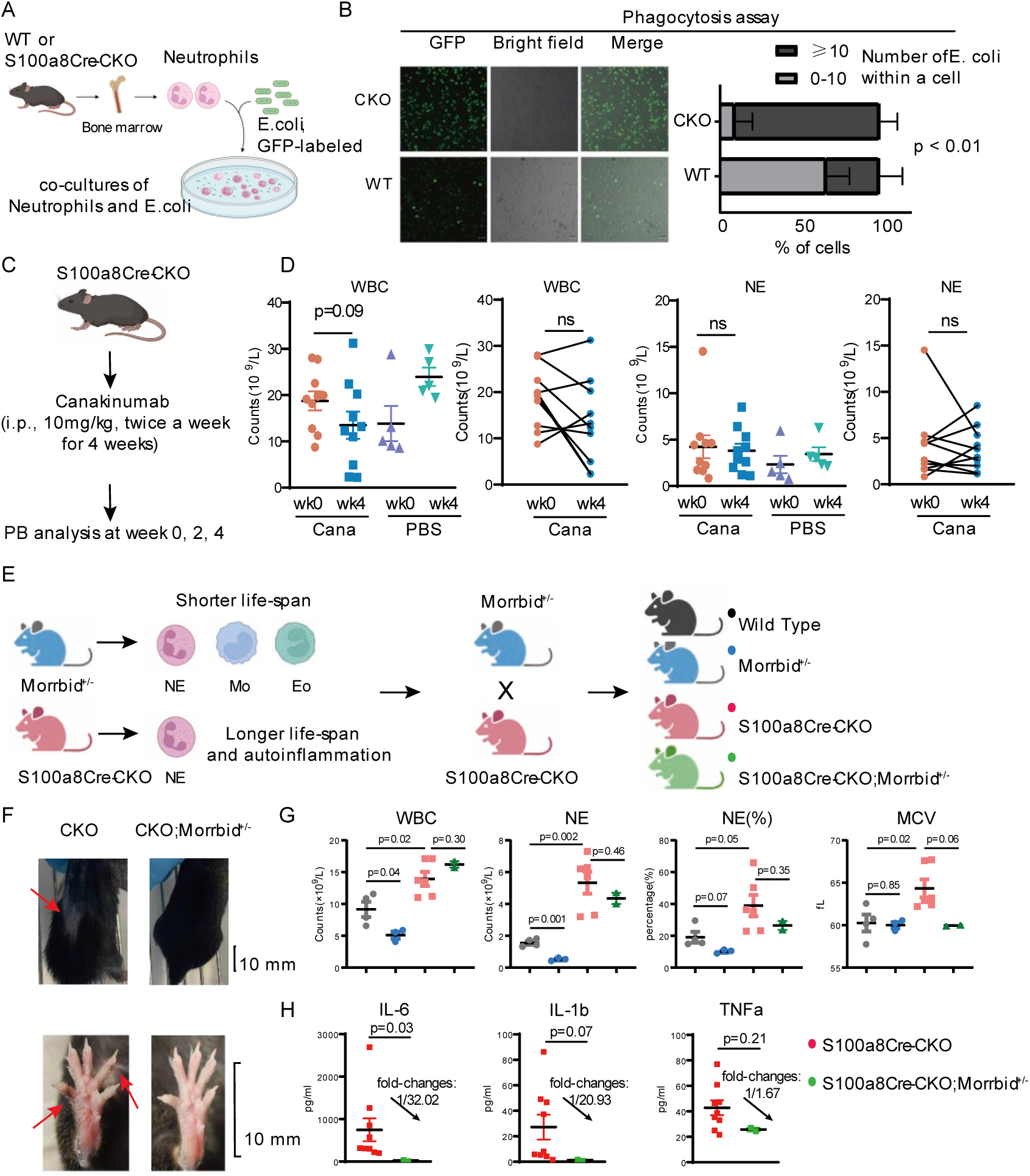
Increased phagocytosis in the mutant neutrophils and perturbation of the VEXAS-like symptoms by drug treatments or by genetic loss of *Morrbid*. (**A-B**) Measuring the phagocytosis capability of the mutant neutrophils. **A**, Scheme for phagocytosis assay. Total BM cells were collected and neutrophils were enriched (see Methods for detail). GFP-labeled *E. coli* was cultured and used for the phagocytosis assays. Once the OD_600_ of GFP-labeled *E. coli* reached to 0.5, *E. coli* was mixed with neutrophils with the ration 10:1 in the culture medium. Time-lapse photography was performed every 10 minutes post the mixing and the GFP dots in the neutrophils were counted at the time point 40 minutes post the mixing (**B**). Mutant neutrophils appear to contain more *E. coli* bacteria (chi-square test, p<0.01). Phagocytosis assay was repeated three times and a representative photography is presented. N=3 biological repeats. (**C-D**) *S100a8Cre-CKO* mice were subjected for Canakinumab treatment. See also **Figure S8** for drug treatment with Anakinra. No pre-tests were performed and the regime of Canakinumab in mice was followed the previous report (56): 10 mg/kg, i.p., twice a week for 4 weeks (Monday and Thursday) (**B**). As shown in **C**, at the end-point of the drug treatment, we observed only reduction of WBC counts while the counts of neutrophils remained unchanged. MCV was also unchanged. (**E**) *S100a8Cre-CKO* mice were bred with *Morrbid^+/−^* for testing the role of *Morrbid* in the autoinflammation disease VEXAS syndrome. The human-mouse conserved lncRNA gene *Morrbid* is a pro-survival regulator for myeloid cells, especially for neutrophils (NE), monocytes (Mo) and eosinophils (Eo). Similar to the homozygous *Morrbid^−/−^*, the heterozygous mutant *Morrbid^+/−^* manifests similar shorter lifespan of the myeloid cells. We generated the compound mutants *S100a8Cre-CKO*; *Morrbid^+/−^*mice along with other controls. (**F**) Hair loss on the back and pigmentation on the toes were mitigated in the compound mutants *S100a8Cre-CKO*; *Morrbid^+/−^* mice, compared to the *S100a8Cre-CKO* mice. (**G**) Hematological analysis on the peripheral blood cells. MCV, NE counts and NE percentage all were with a decreased trend, indicating the mitigation of VEXAS-like symptoms by genetic loss of *Morrbid*. (**H**) Serum level of pro-inflammatory cytokines IL-6, IL-1β, and TNFα all were downregulated in the compound mutants *S100a8Cre-CKO*; *Morrbid^+/−^*mice compared to *S100a8Cre-CKO* mice.

Taken together, results from scRNA-seq datasets, bulk RNA-seq datasets, protein LC/MS datasets and the experimental phagocytosis assays all indicated that depletion of *Uba1* in the non-lethal *S100a8Cre-CKO* mutants result in various aspects of disturbed cellular or immunological activities including UPR, phagocytotic activities and NETs formation, which were also indicated in human VEXAS syndrome patients (17).

### Treatment with IL-1 signaling inhibitors or genetic loss of Morrbid partially mitigated the VEXAS-like symptoms

It is well known that autoinflammatory diseases are generally modulated by IL-1 signaling (47). We tried treatment for the *S100a8Cre-CKO* mutants with Anakinra (a peptide-like antagonist for IL-1R1) or Canakinumab (a monoclonal antibody for IL-1β). As shown in **Figure 6C-D** and **Figure S8**, treatment with Anakinra or Canakinumab only partially mitigated the symptoms in *S100a8Cre-CKO* based on the tested regime: WBC counts were decreased but neutrophil counts appeared unchanged.

In our previous anti-leukemia studies, we demonstrated that the human-mouse conserved lncRNA *Morrbid* is a pro-survival regulator for myeloid cells and has a role in myeloid-lineage leukemogenesis (48) (49) (50) (51). According to the rationale shown in **Figure 6E**, we assessed the role of *Morrbid* by generating the compound mutant *S100a8Cre-CKO;Morrbid^+/−^*. As shown in **Figure 6F-H**, genetic loss of *Morrbid* also partially mitigated the VEXAS-like symptoms including decreased counts of WBC and MCV, disappearance of pigmentation in the toes and decreased level of proinflammatory cytokines.

## DISCUSSION

As illustrated in **Figure 7**, we align the findings from the clinical studies of VEXAS syndrome in human (**Figure 7A**) and that from our study in mice (**Figure 7B**). In the **Supplemental Material Table-S1** to **S4**, we also provide four tables covering the comparison of human VEXAS syndrome and our mouse modeling in details. Overall, it is persuasive that VEXAS syndrome in human is induced by the comprehensive consequence caused by *UBA1* somatic and pathogenic mutations in HSCs while the nine CKO models in the present study recapitulate consequences of *Uba1* loss in certain hematopoietic cell types (a holistic and synthetic effect in human vs. a limited and cell-type specific effect in mouse here). It was suggested that mutant monocytes and mutant neutrophils both contribute VEXAS autoinflammatory symptoms in human. However, when no pathogen or trauma or other challenges had been applied, our study does not support the hypothesis that mutant monocytes with a null mutation of *Uba1* intrinsically play a role in VEXAS-like syndrome in mice. Among the 9 Uba1-CKO mutants described here, only *S100a8Cre-CKO* mice induces VEXAS-like symptoms. In addition to mature monocytes and macrophage*, Lyz2Cre* is reported to have leakage in neutrophils. As *Lyz2Cre;CKO* mice don’t show phenotype as strong as *S100a8Cre;CKO*, Cre activity in neutrophil for *Lyz2Cre* is probably not as strong as that for *S100a8Cre* in the study or the timing of *S100a8Cre* is earlier than that of *Lyz2Cre* for neutrophil depletion of Uba1. Using qRT-PCR and immunological blotting, we experimentally verified that *Lyz2Cre* does have slight leakage in neutrophil but the residual Uba1 protein maintained as high as ∼78%, while residual of Uba1 protein in monocytes is only about 24% (**Figure 3K** and **L**). Lack of abnormalities in *Cx3cr1Cre;CKO* mice provide a further and direct evidence supporting that monocyte or macrophage-loss of *Uba1* is not sufficient to induce VEXAS-like phenotypes in murine models. In addition, fraction of monocytes/macrophages in the BM and PB is much less than that of neutrophils (for example in mouse bone marrow, fraction of monocytes is about 5% while that of neutrophils is about 60%), which represents another reason for lacking of VEXAS-like phenotype in *Lyz2Cre;CKO* and *Cx3cr1Cre;CKO*. Through measuring the gene expression and cell activity by scRNA-seq, we again validated the neutrophil-specific loss of *Uba1* in *S100a8Cre; CKO* and disturbed cellular and immunological activities in various aspects, which are very likely related to the intrinsic function of Uba1 and neutrophils (**Figure 5**).

**Figure 7:**
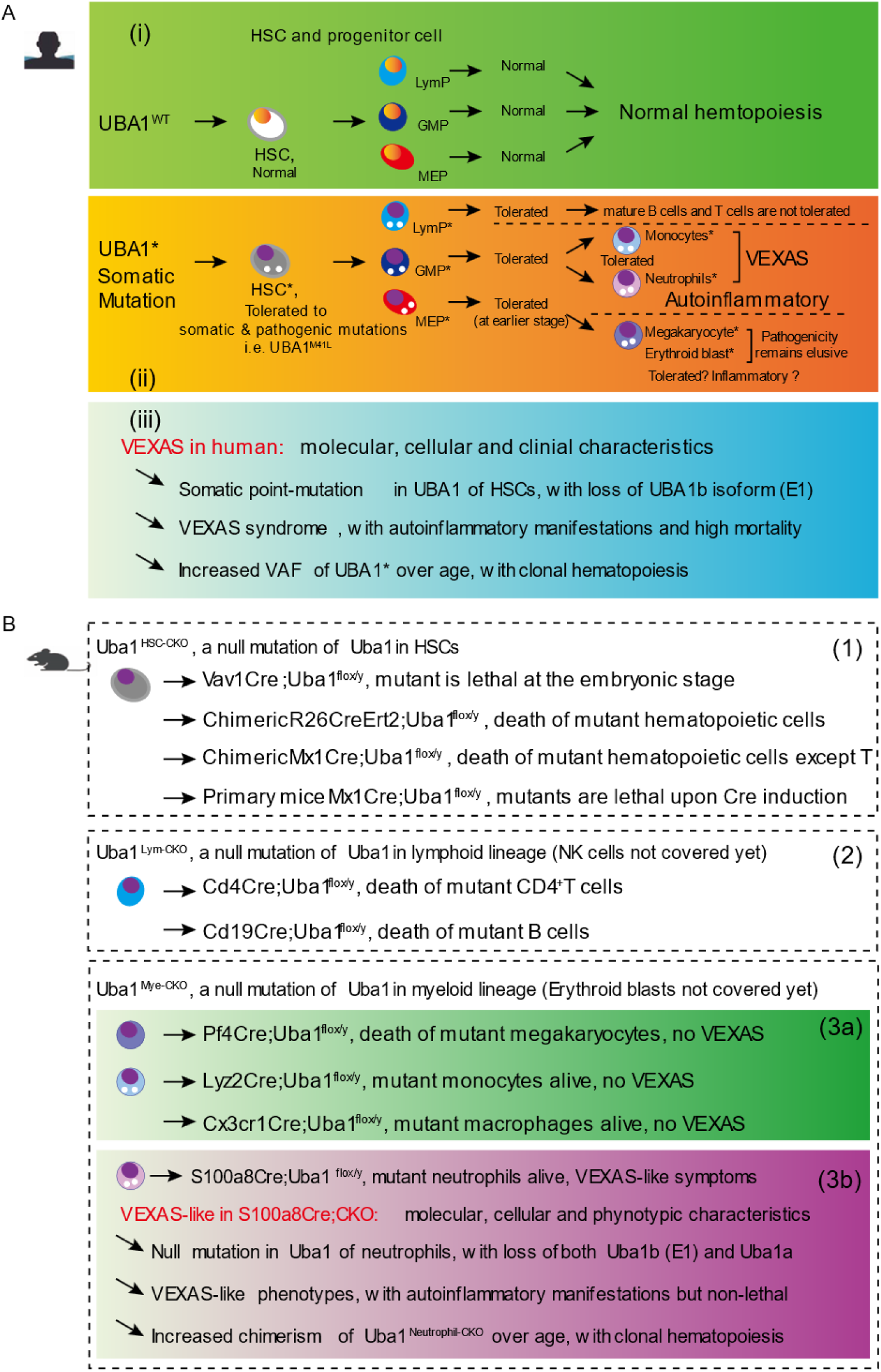
Summary of the study and a comparison between occurrence of VEXAS syndrome in human and VEXAS-like symptoms in *S100a8Cre*;CKO mice. We summarize and compare the occurrence of VEXAS syndrome in human (**A**) and VEXAS-like symptoms in *S100a8Cre*;CKO mice of this study (**B**). See also Discussion in the **Main text** and **Table S1** to **S4** in the **Supplemental Material** for further comparison between current studies in human VEXAS syndrome and studies in the present murine models. **(A)** Normal and VEXAS-conditioned hematopoiesis are illustrated in A-i and A-ii respectively. Additionally, according to the published clinical results, as summarized in A-iii, VEXAS patients manifest three typical features: 1. A somatic and pathogenic point-mutation in *UBA1* of HSCs (typically at the M41 site, indicating a loss of the cytoplasm isoform UBA1b), and VEXAS outcomes should be attributed to the combination effect of all blood cells; 2. Autoinflammatory symptoms at older ages (around 40-year-old to older for aged male); and 3. Increased VAF of mutant *UBA1* and occurrence of clonal hematopoiesis. **(B)** The 9 different CKO lines of the study are summarized in B-1, B-2, B-3a and B-3b respectively. We demonstrated that most of hematopoietic cells are not tolerated to the null-version mutation of *Uba1* while monocytes and neutrophils survive with ∼30% residual level of Uba1. As summarized in B-3b, VEXAS-like symptoms in *S100a8Cre*;CKO also include three typical features: 1. A null-version of mutation in *Uba1* of neutrophils, indicating both nucleic and cytoplasm isoforms are deficient and consequences are mainly attributed to mutant neutrophils; 2. Autoinflammatory symptoms typically start to occur at 3-month-old and extend to older ages (i.e. 6-month-old to older) but its life-span are grossly normal; and 3. Increased fraction of mutant neutrophils and occurrence of clonal hematopoiesis as indicated by the cBMT experiments.

We are aware that *S100a8Cre-CKO* mice are indeed not as the same as the point mutations in human: a null version of mutation vs. a missense version of mutation, but interestingly both are loss-of-function, which is one of the key etiologies of VEXAS syndrome. Reversing such loss-of-function defects could guide future development of interventions for this important biological process mediated by UBA1. When muting the function of Uba1 in almost of each major cell type in the bone marrow, our study demonstrates that null-mutations of *Uba1* in neutrophils rather than in other hematopoietic cells are critical for inducing VEXAS-like symptoms. Nonetheless, installing a point-mutation in mouse HSCs by gene editing is undoubtedly necessary and demanded to fully recapitulate VEXAS symptoms in mice, which is suggested by *Molteni et al* in a recent meeting abstract (35).

Human UBA1 and mouse Uba1 are highly conserved and critical for eukaryotic cells. A quick and extensive death of hematopoietic cells took place in the HSC-wise depletion of *Uba1* (*Vav1Cre*, *R26CreErt2* and *Mx1Cre* mediated respectively) suggesting an HSC with a null mutation of Uba1 (*Uba1^null^*) is not tolerated at all. In contrast, tolerance of somatic mutations in human HSC (i.e. *UBA1^M41L^*) is quite intriguing and suggests further functional studies are required to ask if compensation function of UBA1a isoform or other compensation mechanisms (i.e. UBA6) exist in human HSC with *UBA1^M41L^* but those mechanisms does not take place in mouse HSC with *Uba1^null^*. Of note, no VEXAS-like phenotypes were observed in the two HSCs-targeting chimeric mice with various chimeric fractions as analyzed in cBMT assays using *Mx1Cre;*CKO or *R26CreErt2;*CKO donor cells. These results further suggest that an HSC with *UBA1^M41L^* is <u>unequal</u> to HSC with *UBA1^null^*. “Tolerated or not tolerated” and “pathogenic or not pathogenic”, will represent two key questions in future to ask each individual variant among the full spectrum of mutations identified in the available VEXAS syndrome cohorts.

Although we provided data with regards to the intolerance of megakaryocytes by *Pf4Cre;* CKO, the cell-type-dependent tolerance and pathogenicity of erythroid blast and megakaryoblast, among other intermediate myeloid and lymphoid progenitors and blast cells, will still be one of the interesting questions to ask in future (i.e. experimental and functional examination). Using VEXAS patient cells for a direct test under the *ex vivo* or *in vitro* experimental settings are demanded. In addition, inducible *Cre* lines targeting hematopoietic progenitors (i.e GMP, MEP and lymphoid progenitor) rather than HSCs at the top of hematopoietic hierarchy will be worthwhile too to dissect the role of Uba1 along the full journey and each branch of hematopoiesis.

In contrast to intolerance of *Uba1^null^* for HSCs, we uniformly observed the tolerance of *Uba1^null^* for monocytes and neutrophils, which were also observed in human mutant monocytes and neutrophils with *UBA1^M41L^*. Although flow cytometry-based apoptosis profile of mutant neutrophils suggests a higher apoptotic activity, we provided couples of alternative *in vivo* evidences supporting the survival advantage of mutant neutrophils when comparing with the normal neutrophils: (1) hematological cell counts in the peripheral blood, (2) cBMT assays, (3) half-life tracing by BrdU staining, and (4) lox-GFP-based genetic tracing. We speculate that the increased expression of pro-inflammatory cytokines IL-6, IL-1β and TNFα is the driver of the survival advantage of the mutant neutrophils and a positive feedback loop appears to take place in the VEXAS disease progression. Such a positive-feedback loop model has been proposed in our previous studies about *Tet2* mutation and clonal hematopoiesis (48,49,50,51). It would be interesting to test in clinic in future if mitigating clonal hematopoiesis and such positive-feedback loop brings any benefits for VEXAS syndrome patients.

Since the human VEXAS syndrome is a synthetic effect of all aspects of mutant hematopoietic cells carrying somatic *UBA1* mutations, it therefore will be required in future to generate compound *Cre* lines to test if stronger phenotype take place (i.e. generating *Lyz2Cre; S100a8Cre; Uba1^flox/y^*). Furthermore, Anakinra, an inhibitor of IL-1/IL-1R1 signaling, and Ruxolitinib, an inhibitor of JAK2 signaling, have been reported to treat VEXAS syndrome (52–53). It will be interesting to test if combination of these two drugs alleviate the autoinflammatory symptom in *S100a8Cre-CKO* mice. In addition, in our recently submitted work on *Pstpip2*-KO induced autoinflammatory disease chronic multifocal osteomyelitis (CMO) and in *Nox2*-KO induced chronic granulomatous disease (CGD), we demonstrated that genetic loss of *Morrbid* mitigated these two autoinflammatory diseases too (Preprints, 57, 58). In addition to its function in anti-leukemia, targeting *Morrbid* may represent a valuable strategy for anti-inflammation in future clinical management.

### Limitation of the study

In the present study we introduced a null version of mutation in *Uba1* rather than a point mutation like *Uba1^M41L^* in murine models. Furthermore, Multiple *Uba1^non-M41^* variant were reported. However, we should be cautious that the pathogenicity of those non-M41 variant require functional experiment to validate (54). Our study provides an indirect but complementary and timely understanding of VEXAS syndrome. In addition, the present study has not covered the respective roles of *Uba1* for erythroid blast, mast cells, eosinophils and basophils but all other major cell types were covered. To fully dissect the role of *Uba1* in hematopoietic system, additional *Cre* lines are required to breed with the *Uba1* flox knockin line. Furthermore, our preliminary phagocytosis experiments indicated that the mutant neutrophils appear to have stronger NETs and phagocytosis capabilities. Given that the easy maintenance of the *S100a8Cre; Uba1^flox/y^* colonies, it will be interesting to test the mutant mice manifest any aberrations in pathogen-host immunity, trauma, cancer immunity, and other pathophysiological challenges.

## CONCLUSION

In conclusion, we report a VEXAS-like mouse model *S100a8Cre;Uba1^flox/y^*for a generally ignored autoinflammatory disease in human: the VEXAS syndrome. Using the other eight CKO models, we also demonstrate that targeting some hematopoietic cell types, rather than neutrophils, failed to manifest VEXAS-like symptoms. These results well document the intrinsic and diversified functions of Uba1 in mammal hematopoietic cells and suggest cell-type-dependent pathogenicity of *UBA1^mut^* in human VEXAS syndrome. The VEXAS-like mouse model *S100a8Cre-CKO* will assist further understanding of VEXAS syndrome and warrant future development of treatments for the newly identified autoinflammatory disease highly prevalent among aged men.

## METHODS

### Mice

As the VEXAS syndrome is a disorder mostly affecting aged men, we mainly generated male CKO mutant mice (*Cre; Uba1^flox/y^*) and collected the tissue samples from the male mutant mice for the following studies described below.

The *Uba1^flox/y^* mice were generated by the CRISPR/Cas9 approach. The Cre mice were described previously and purchased from Jax Laboratory. The dTomato-lox-GFP mice (named here as lox-GFP for simplicity) were from Jax Laboratory. The Cre and flox strained used in the study all are in C57/B6 background. Except *Vav1Cre*-CKO, we used other CKO male mice at the age of 8 to 24 weeks and the *Uba1^flox/y^*or corresponding *Cre*-positive males in the same litter or with same ages as controls. Animal experimentation was performed in accordance with protocols approved by the Animal Care and Use Committee of Tianjin Medical University.

### H&E Staining

The tissue samples were fixed in 4% paraformaldehyde. After the dehydration in ethanol, the tissue was embedded in paraffin then sectioned with a thickness of 4 μm. The histopathological feature was tested via H&E staining.

### Flow cytometry

The bone marrow (BM) cells and peripheral blood mononuclear cells (PBMCs) were flushed out with FACS buffer. Single-cell suspensions were treated with red blood cell lysis buffer, stained, and analyzed using FACS Canto II (BD Biosciences). For mature cell analysis, staining with antibodies against CD19 (#152410, BioLegend), CD3 (#100206, BioLegend), Gr-1 (#108408, BioLegend), CD11b (#101206, BioLegend), CD4 (#100510, BioLegend), CD8a (#100712, BioLegend), CD41 (#133904, BioLegend). For early hematopoietic cell analysis, cells were incubated with biotinylated antibodies against the lineage (Lin) markers: CD11b, Ter119 (#116208, BioLegend), Gr-1, CD3, B220, and the fluorescence-conjugated antibodies: c-Kit (#105812, BioLegend), Sca-1 (#108126, BioLegend), CD34 (#11-0341-85, BioLegend), CD150 (#115912, BioLegend), CD48 (#103424, BioLegend), and CD16/32 (#101318, BioLegend).

### Competitive bone marrow transplantation (cBMT) and chimerism assays

Recipient animals (BoyJ, CD45.1) were lethally irradiated (7Gy plus 4Gy) one day prior to transplantation (intravenous tail injection) of donor cells. For generating chimeric mice mimicking hematopoietic clonal expansion, *Cre;Uba1^flox/y^* donor cells and F1 donor cells were mixed at a ratio of 1: 1 (100K:100K). 4 weeks after BM *R26CreErt2;Uba1^flox/y^*transplantation animals were continuously administered by gavage with tamoxifen at a dose of 15mg/ml in 200ul of corn oil for 5 days. *Mx1Cre;Uba1^flox/y^* was injected with PolyI:C at a dose of 1.5mg/ml in 100ul of PBS by intraperitoneal injection three times on alternate days.

### Neutrophil phagocytosis assays

Bone marrow cells were isolated from age and sex matched mice (12-week old), followed by neutrophil enrichment subsequently using the density gradient centrifugation (Sigma-Aldrich Cat. No.11191, No.10771) according to the manufacturer’s instructions. The enriched neutrophil sample purity was assessed by flow cytometry with CD11b and Ly6G, confirming expected purity. GFP-labeled *E. coli* bacteria were grown overnight to an OD600 of 0.5, and subsequently co-cultured with neutrophils. For co-culture, 5×105 cells in 0.5mL of phenol red-free RPMI (Gibco) supplemented with 10% pooled mouse serum were plated per 35mm glass bottom imaging dish (Mattek corporation) and mixed with GFP-labeled *E. coli* bacteria at the ratio 1:10. Time lapse imaging was performed using the confocal scanning microscope (Leica) and six fields of view were captured over 40 min per sample. Phagocytosis was analyzed in FIJI software after stitching the six fields of view together.

### Enzyme linked immunosorbent assay (ELISA)

ELISA detection for ten inflammatory factors (IFN-γ, IL-1β, IL-2, IL-4, IL-5, IL-6, KC/GRO, IL-10, IL-12p70, TNF-α) were performed using Proinflammatory Panel-1 (mouse) Kits (MSD, NJ, USA). Mouse blood samples were collected using an empty tube without anticoagulant, left to stand at 4°C for 2 hours (to avoid shaking and prevent hemolysis), centrifuged for 20 minutes (4°C, 3000g), and collected the supernatant. The samples were frozen at −80°C. Before the experiment, the reagent was moved to room temperature for 30min. According to the kit instructions, we added 25-50uL of standard, quality control, and sample sequentially at room temperature (700rpm, 1h). Then we added 25-50uL of detection antibody and incubate at room temperature (700rpm, 1h). The samples underwent 3 washes with 150uL cleaning solution, followed by the addition of 150uL of plate reading solution before machine analysis. Analysis was carried out using MSD Discovery Workbench (Version 4.0).

### BrdU pulse-phase assay for measuring half-life of neutrophils

Mice were injected intraperitoneally with 2 mg of BrdU (Sigma-Aldrich) at the indicated time points (Day −8 to Day −1, one dose each day prior to the BrdU flow cytometry assays). To detect BrdU incorporation, cells from the blood were stained with neutrophil surface makers CD11b and Ly6G. Cells were then fixed, permeabilized, and finally stained intracellularly with APC-conjugated anti-BrdU antibody according to the manufacturer’s protocol before analysis with flow cytometry.

### Drug Treatments for the mutants

Anakinra (recombinant interleukin-1 receptor IL-1R1 antagonist): the *S100a8Cre-CKO* mice were treated by i.p. injection with Anakinra (37mg/kg) or PBS for four times a week. Blood examination was recorded at Week 0, 2, and 4 post the treatment. Canakinumab (recombinant IL-1β monoclonal antibody): the S100a8Cre-CKO mice were treated by i.p. injection with Canakinumab (10mg/kg) or PBS for twice a week. Blood examination was recorded at Week 0, 2, and 4. At Week 4 post the treatment, flow cytometry was used to detect neutrophils in the blood.

### Total protein extraction

The samples were taken out in the frozen state and put on ice. The samples were suspended in protein lysis buffer (8M urea,1% SDS) which included appropriate protease inhibitor to inhibit protease activity and the mixture were treated by high-flux tissue grinding machine for 3 times, 40 s each. Then the mixture was incubated on ice for 30 min, during which was vortex mixed for 5-10 s every 5 min. After centrifugation at 16000g at 4°C for 30 min, the concentration of protein from the supernatant collected were determined by Bicinchoninic acid (BCA) method by BCA Protein Assay Kit (Thermo Scientific). Protein quantification was performed according to the kit protocol.

### Western blot analysis

Proteins were harvested from BM cells with RIPA Lysis Buffer (Beyotime, Jiangsu, China) supplemented with phenylmethyl sulfonyl fluoride (PMSF) protease inhibitor and phosphatase inhibitor. Total protein concentration was determined by BCA Protein Assay Kit (Beyotime, Jiangsu, China), denatured protein samples of appropriate quality of proteins were subjected to sodium dodecyl sulfate polyacrylamide gel electrophoresis (SDS-PAGE) and then transferred to PVDF membranes. Then membranes were later blocked with 5% skimmed milk, and incubated were immunodetected with specific antibodies against UBA1 (#4891; Cell signaling technology, USA), antibodies against Uba1a (#4890; Cell signaling technology, USA), antibodies against UBA6 (#13386; Cell signaling technology, USA), antibodies against Free Ubiquitin and Polyubiquitin (#43124; Cell signaling technology, USA), antibodies against Ubiquitin (#3936; Cell signaling technology, USA), Actin (AC026, ABclonal, China) and GAPDH (#ab181620, Abcam, USA) overnight at 4 °C. Protein bands were visualized by the MINICHEMI Imaging System (Surwit, Hangzhou, China) using the commercial Pierce™ Fast Western Blot Kit and the ECL Substrate (GenStar, Beijing, China).

### Bulk RNA-Seq Data Analysis

RNA-seq analysis pipeline was conducted in house and has been described in our previous studies (55). Volcano plots were used to show the fold changes and log-adjusted p-values for DEGs. We used the R package “clusterProfiler” to perform Gene Ontology (GO) analyses. “enrichGO” and genome-wide annotation packages “org.Mm.eg.db” were needed. A p-value < 0.05 was considered significant enrichment.

### Single-cell RNA-sequencing (scRNA-seq) analysis

#### data acquiring and processing

Single-cell RNA-Seq libraries were prepared using SeekOne® MM Single Cell 3’ library preparation kit (No. K00104, SeekGene). Briefly, the appropriate number of cells were loaded into the flow channel of SeekOne® MM chip which had 170,000 microwells and allowed to settle in microwells by gravity. After removing the unsettled cells, sufficient Cell Barcoded Magnetic Beads (CBBs) were pipetted into flow channel and also allowed to settle in microwells with the help of a magnetic field. Next excess CBBs were rinsed out and cells in MM chip were lysed to release RNA which was captured by the CBB in the same microwell. Then all CBBs were collected and reverse transcription were performed at 37℃ for 30 minutes to label cDNA with cell barcode on the beads. Further Exonuclease I treatment were performed to remove unused primer on CBBs. Subsequently, barcoded cDNA on the CBBs was hybridized with random primer which had reads 2 SeqPrimer sequence on the 5’ end and could extend to form the second strand DNA with cell barcode on the 3’ end. The resulting second strand DNA were denatured off the CBBs, purified and amplified in PCR reaction. The amplified cDNA product was then cleaned to remove unwanted fragments and added to full length sequencing adapter and sample index by indexed PCR. The indexed sequencing libraries were cleanup with SPRI beads, quantified by quantitative PCR (KK4824, KAPA Biosystems) and then sequenced on illumina NovaSeq 6000 with PE150 read length. The raw gene expression matrices of the scRNA-seq data were read and combined, converting them into a Seurat object using the Seurat package (version 4.4.0) in R software (version 4.2.0). During the quality control process, we retained cells meeting the following criteria: nFeature_RNA >200 and nFeature_RNA < 5000; and UMIs derived from the mitochondrial genome < 5. We excluded genes related to mitochondria from the datasets. We performed the function FindVariableFeatures (Seurat) to detect the features with highest coefficient of variation (CV). By default, we chose the top 2000 variable features to calculate a PCA matrix with 30 components and transported the PCA matrix into Harmony (version 1.2.0) to integrate single-cell data gene expression matrix and correct the batch effect.

#### unsupervised clustering and annotation of cell types in the UMAP plots

The Harmony matrix would be used for unsupervised clustering by building the nearest neighbor graph and Louvain algorithm. The unsupervised clustering (resolution = 0.3) for identifying the main cell types, including the HSPCs (Ms4a3, Cebpd), Pro Neutrophils (Camp, Chil3), Neutrophils (Mmp8), Monocytes (F13a1, Ms4a6c), Erythrocytes (Hba-a2, Hbb-bt), pDC (Siglech, Cox6a2), Basophils (Prss34, Mcpt8), B (Cd19), Pro B (Vpreb3), T (Cd3e), Plasma cell (Jchain, Iglc2).

#### construction of gene signatures and scoring bioactivities in each cell

We integrated classic gene sets to characterize different bioprocesses, generating gene lists for subsequent analyses. The function ‘AddMouduleScore’ in the Seurat package was employed to calculate the average expression levels for each cluster. All signatures were binned based on the average expression.

### Proteomic analysis

#### protein reductive alkylation and digestion

Take protein samples 100 μg and add TEAB (Triethylammonium bicarbonate buffer), which the final concentration of TEAB is 100 mM. Then add TCEP (tris (2-carboxyethyl) phosphine) to the final concentration of 10 mM and react for 60 min at 37 °C. Following add IAM (Iodoacetamide) to the final concentration of 40 mM and react for 40 min at room temperature under dark conditions. Add a certain percentage (acetone: sample v/v = 6:1) of pre-cooled acetone to each sample and to settle for 4 h at −20 °C. After centrifugal for 20 min at 10000 g, the sediment was collected and add 100 µL 100mM TEAB solution to dissolve. Finally, the mixture was digested with Trypsin overnight at 37 °C added at 1:50 trypsin-to-protein mass ratio.

#### protein extraction and quantification

Samples were thawed on ice and processed using a high-throughput tissue grinder in a lysis buffer containing 8M urea and 1% SDS to ensure thorough lysis. Additionally, to prevent proteolytic degradation during the process, a cocktail of protease inhibitors was added. Samples were homogenized in three cycles of 40 seconds each, followed by a 30-minute incubation on ice with vertexing every 5 minutes to enhance protein extraction. The mixture was then centrifuged at 16,000g for 30 minutes at 4°C to separate the soluble proteins from cellular debris. The supernatant was carefully collected, and protein concentration was determined using the Bicinchoninic Acid (BCA) Protein Assay Kit (Thermo Scientific), following the manufacturer’s instructions to ensure accurate quantification.

#### protein digestion

Proteins were diluted to a final concentration of 100 μg in 100 mM Triethylammonium bicarbonate (TEAB) and reduced with 10 mM Tris(2-carboxyethyl) phosphine (TCEP) at 37°C for one hour. Subsequent alkylation was performed with 40 mM iodoacetamide (IAM) in the dark at room temperature for 40 minutes to protect sensitive amino acid residues. Proteins were then precipitated using cold acetone at a ratio of 6:1 (v/v), incubated at −20°C for 4 hours to enhance precipitation, and centrifuged. The resulting pellet was resuspended in 100 mM TEAB for enzymatic digestion. Trypsin was added at a 1:50 enzyme-to-protein ratio and the mixture was incubated overnight at 37°C to achieve complete digestion.

#### peptide processing and desalting

Following digestion, peptides were vacuum dried and resolubilized in 0.1% trifluoroacetic acid (TFA) for desalting. This step was critical to remove salts and other contaminants that could interfere with subsequent mass spectrometric analysis. Desalting was performed using HLB cartridges, and peptides were eluted, collected, and again subjected to vacuum drying. Peptide quantification was performed using a specialized Peptide Quantification Kit from Thermo Fisher Scientific, following the provided protocol to ensure consistent results.

#### phospho-peptide and ubiquitylated peptide enrichment (UPE)

For selective enrichment of phospho-peptides, the peptide was first dissolved using Binding/Wash Buffer, adding a Binding/Wash Buffer equilibrated column, centrifugation, and then the redissolved peptide was added to the column and incubated for 30min. After incubation, add Binding/Wash Buffer cleaning column, centrifuge; add LC_MS grade water to clean the column, then add Elution Buffer stripping twice; finally, add equal volume of 20% TFA acidification, stage-tip desalination, and drain by vacuum concentrator. For ubiquitylated peptides, enrichment was carried out using Anti-K-ε-GG antibody beads provided by Cell Signal Technology, which specifically recognize diglycine-modified lysine residues. Both types of peptides were bound to their respective beads under optimized conditions, washed to remove non-specifically bound peptides, and the bound peptides were eluted. The eluted peptides were then desalted using C18 Stage Tips to prepare them for LC-MS/MS analysis.

#### LC-MS/MS analysis

Peptide mixtures were analyzed using a state-of-the-art VanquishNeo system coupled to an Orbitrap Astral mass spectrometer (Thermo, USA). Separation was achieved on a C18-reversed phase column with a gradient of acetonitrile in 0.1% formic acid. Peptides were analyzed in data-independent acquisition (DIA) mode, which allows for a comprehensive survey of all peptides present, with a mass range set from m/z 100 to 1700.

#### data analysis

Raw data files were processed using the Spectronaut software (Version 18), with a stringent false discovery rate (FDR) of ≤0.01 for protein and peptide identifications to ensure high confidence in the results. Quantitative analysis was based on the integrated peak areas of at least six peptides per protein and three daughter ions per peptide, providing robust quantification metrics.

#### bioinformatics analysis

Differentially expressed or modified proteins were identified using the Majorbio Cloud platform, with statistical significance set at p < 0.05 and fold change thresholds. Functional annotations and pathway analyses were conducted using the Gene Ontology (GO) and Kyoto Encyclopedia of Genes and Genomes (KEGG) databases to elucidate biological implications of the proteomic changes observed.

### Statistical analysis

Statistical analysis was conducted by using GraphPad 9. Comparisons between 2 groups were determined by using a two-tail *Student’s* t-test or by using *Wilcoxon* test. Comparison of multiple groups were determined by using an ANOVA analysis of variance with the Dunnett multiple comparisons test. Results with p < 0.05 were considered statistically significant.

## Author contributions

ZC is the guarantor of the study. ZC conceived, designed, supervised the study, analyzed the data and wrote the manuscript. GD, JL, HZ, WJ, YW, and ZW performed the experiments, analyzed data, edited the manuscript, and contributed equally to the project. KX, JZ, LM, YM, LCC, QZ, HW, WW, and YF assisted the experiments, analyzed the data and edited the manuscript.

## Acknowledgements

We thank our colleagues for technical support, critically reading our manuscript, and their suggestions to improve the manuscript. We would also like to thank Drs. Liu, Li, Ding, Zhang and Yu (Tianjin Medical University School of Basic Medicine), and Cheng (Chinese Academy of Medical Sciences, National Clinical Research Center for Blood Diseases) for sharing mice.

## Ethical approval information

Animal experimentation was performed in accordance with protocols approved by the Animal Care and Use Committee of Tianjin Medical University.

## Data sharing statement

Raw bulk RNA-seq and scRNA-seq data in this study has been deposited in China National Center for Bioinformation/Beijing Institute of Genomics, Chinese Academy of Sciences: https://ngdc.cncb.ac.cn/gsa/. (GSA: CRA105841). Proteomic data and scRNA-seq Matrix and have been deposited in https://ngdc.cncb.ac.cn/omix/. (OMIX006177& OMIX006178). Omics data will be available upon publication.

## Patient and Public Involvement

Not Applicable.

## LEGENDS for Supplemental Figures

**Figure S1:**
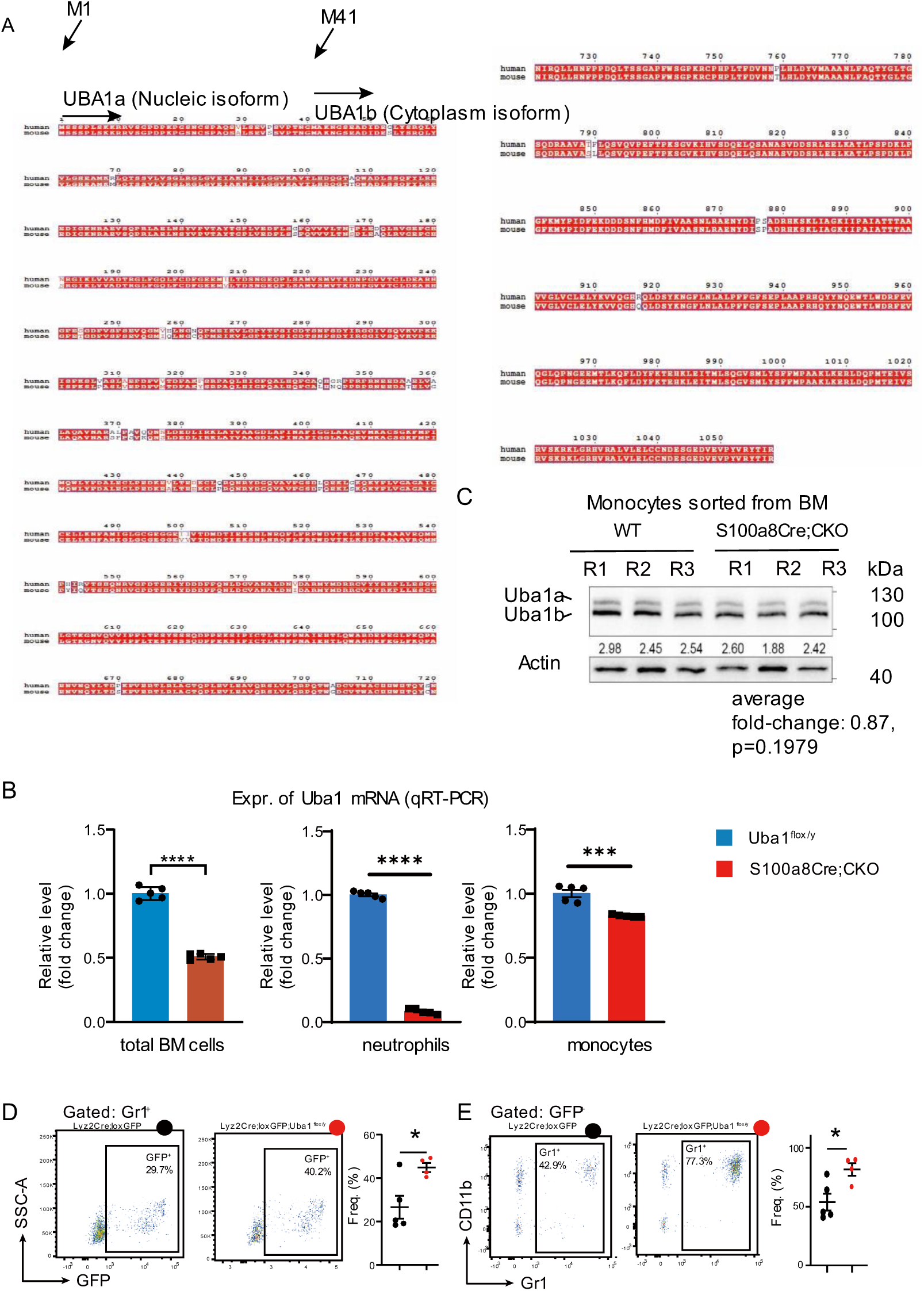
Conservation of UBA1 protein sequences in human and mouse and experimental assays on depletion efficiency of Uba1 in the CKO models. Related to **Figure 1** and **3** in the **Main text**. **(A)** Protein sequences of human UBA1 and mouse Uba1 are compared. Mouse and human express UBA1 in same length of BM cells (1058 aa) with >95% sequence identity. The start point of UBA1 isoform is at M1 while that of UBA1b isoform is at M41. Related to Figure 1. **(B)** Quantification of *Uba1* mRNA expression in total BM cells, neutrophils and monocytes of S100a8Cre;CKO by qRT-PCR. Box plots show the relative expression level of *Uba1* in total BM cells, purified neutrophils and monocytes. Related to Figure 1. **(C)** Quantification of Uba1 protein expression in monocytes of S100a8Cre;CKO by western blotting. Related to Figure 1. **(D)** *Lyz2Cre; lox-GFP* and *Lyz2Cre; lox-GFP; Uba1^flox/y^* mice were generated and used for confirmation of *Lyz2Cre* activity in the study. We crossed the *Lyz2Cre; Uba1^flox/y^* CKO mutants with tomato-lox-GFP line to generate *Lyz2Cre; lox-GFP* and *Lyz2Cre; lox-GFP; Uba1^flox/y^*. Expression of GFP indicates the activity of *LyzCre* in the Cre/loxP strategy. When gated with Gr1^+^ channel or with GFP^+^ channel, we confirmed that *Lyz2Cre* activity is robust (an indirect indication for loss of *Uba1*) in Gr1^+^ myeloid cells in *Lyz2Cre; lox-GFP; Uba1^flox/y^*. Interestingly such loss of *Uba1* in *Lyz2Cre; lox-GFP; Uba1^flox/y^* did not induce death in Gr1^+^ myeloid cells. In contrast, mutant Gr1^+^ myeloid cells appear to have longer life-span than that WT Gr1^+^ myeloid cells. Related to Figure 3. ns, not significant; *, p<0.05; **, p<0.01; ***, p<0.001; ****, p<0.0001; n=3∼5 biological repeats.

**Figure S2:**
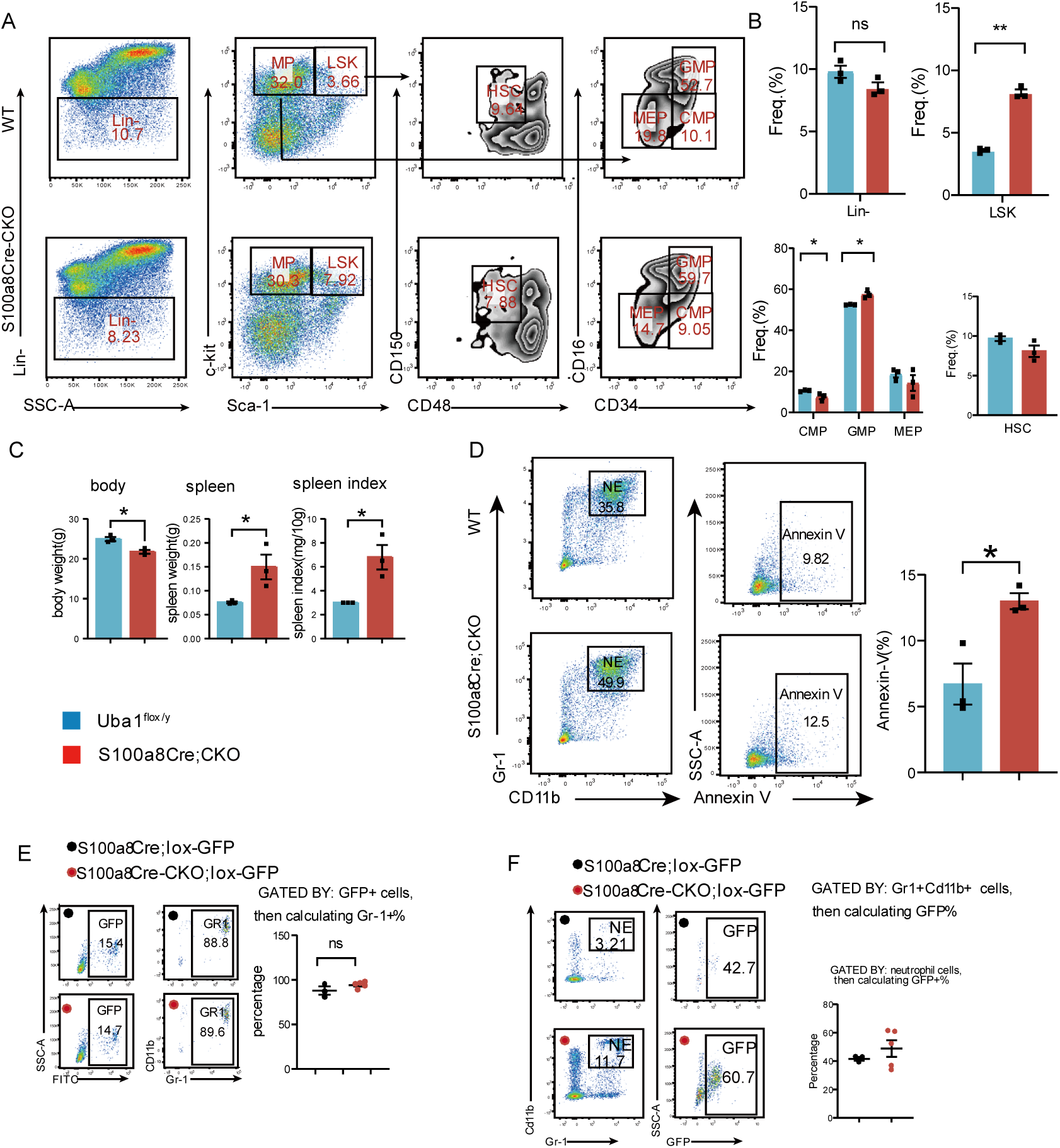
Flow cytometry analysis on bone marrow cells from *S100a8Cre-CKO* mice. Related to **Figure 4** in the **Main text**. Potential hematopoietic abnormalities were examined by flow cytometry using bone marrow (BM) or peripheral blood cells. BM or PB cells from at least 3 WT animals and 3 S100a8Cre-CKO animals for the flow cytometry examination. **(A)** Gating strategies and representative flow cytometry profiles for hematopoiesis using bone marrow cells from WT and *S100a8Cre-CKO* mice. **(B)** Quantification of various compartment of hematopoietic progenitor cells as indicated. **(C)** Body weight, spleen weight and spleen index in the WT and S100a8Cre;CKO mice. **(D)** Apoptosis analysis of neutrophils in the bone marrow. Left panel, Gating strategies and representative flow cytometry profile; Right panel, Quantification of apoptotic cells (Annexin-V^+^) in neutrophils. (**E-F**) *S100a8Cre-CKO* mice were bred with lox-GFP mice for validating the activity of S100a8Cre. **E**, GFP^+^ cells were gated and for calculating myeloid cells. **F**, neutrophil cells were gated and for GFP^+^ cells.

**Figure S3:**
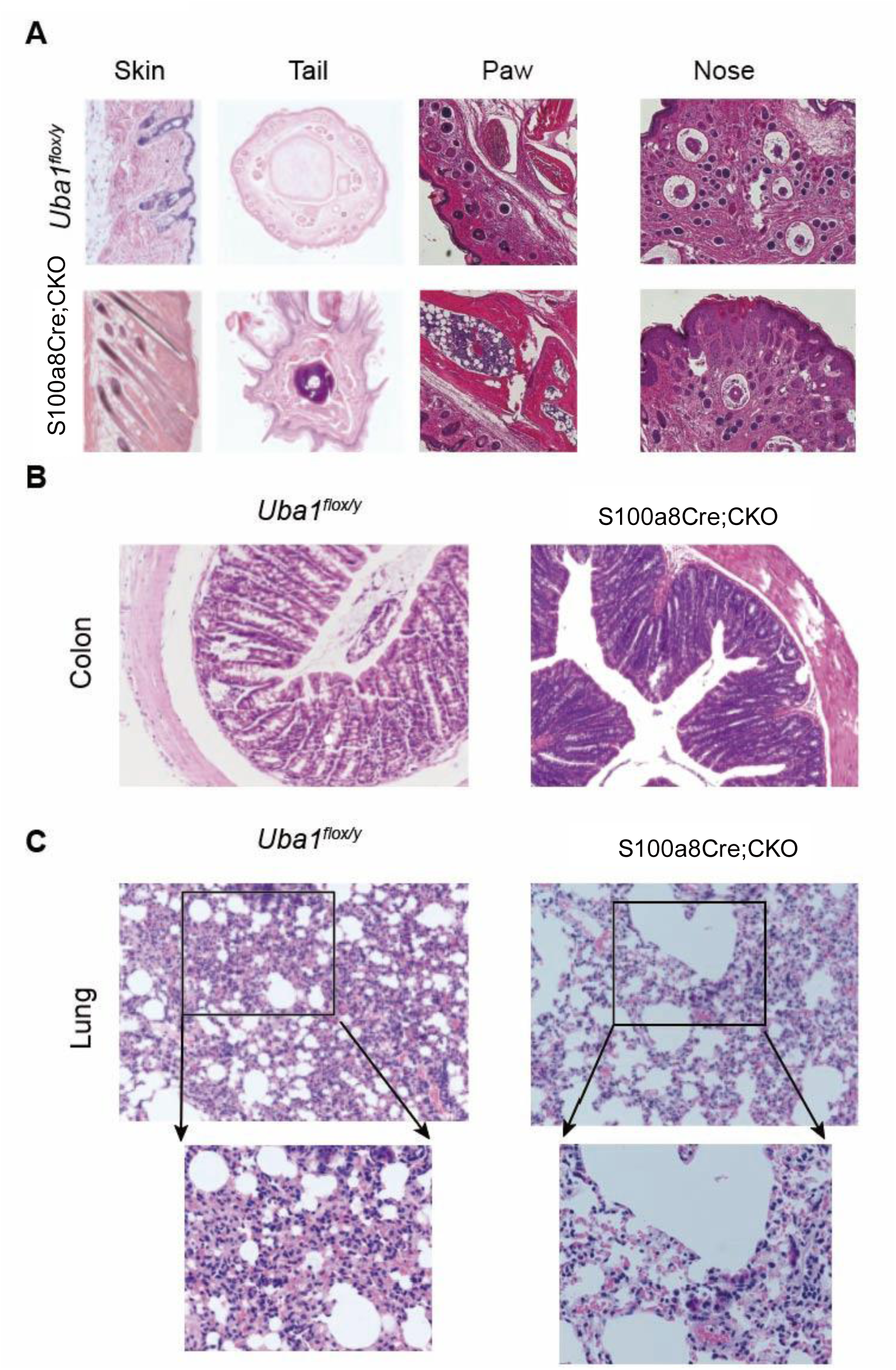
H&E staining of various organs from *S100a8Cre-CKO* mice. Related to **Figure 4** in the **Main text**. Affected tissues were collected from at least 3 WT animals and 3 S100a8Cre-CKO animals for H&E histology examination. **(A)** H&E staining of affected tissues including skin, tail, paw and nose. Note that skin at the back and tail in *S100a8Cre-CKO* mice is not as smooth as in WT. Infiltration of immune cells are observed in the swollen paws in the *S100a8Cre-CKO* mice. Although flare noses in the *S100a8Cre-CKO* mice were observed, we failed to observe much dramatic difference in the H&E histology. **(B)** H&E staining of colon tissues. **(C)** H&E staining of lung tissues.

**Figure S4:**
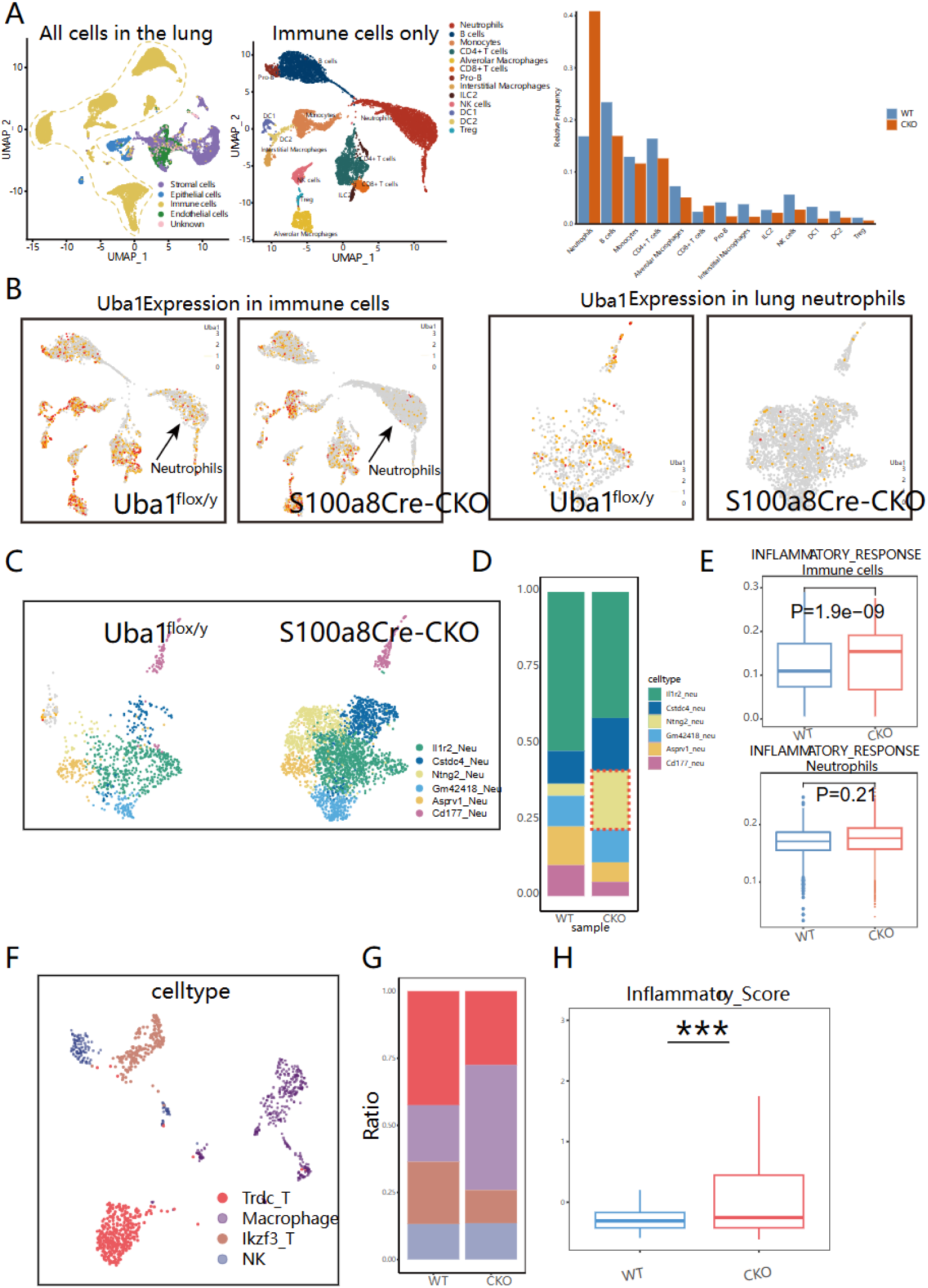
scRNA-seq analysis of lung tissues from *S100a8Cre-CKO* mice. Relying on scRNA-seq dataset, we observed an obvious increased inflammation in the lung tissue from *S100a8Cre-CKO* mice. Tissues were from 1 WT animal and 1 *S100a8Cre-CKO* animal. Related to **Figure 4** in the **Main text**. (**A**) Lung tissues from WT and *S100a8Cre-CKO* mice were subjected for scRNA-sequencing analysis for identifying any possible pathophysiological alterations in the *S100a8Cre-CKO* mice. Immune cells were extracted and subject for further analysis. Left panel: the UMAP plot of total cells of the lung tissue; middle panel: the UMAP plot of the immune cells of the lung tissue; right panel: proportions of various immune cell types in the immune cell pool. (**B**) Expression of *Uba1* in the immune cell pool and in the extracted neutrophils. (**C-D**) Sub-clusters of the neutrophils and their proportions. (**E**) Inflammatory scoring of the total immune cells or just neutrophils (**F-H**) Analysis of other immune cells except neutrophils by re-clustering(**F**), proportion analysis (**G**) and scoring (**H**).

**Figure S5:**
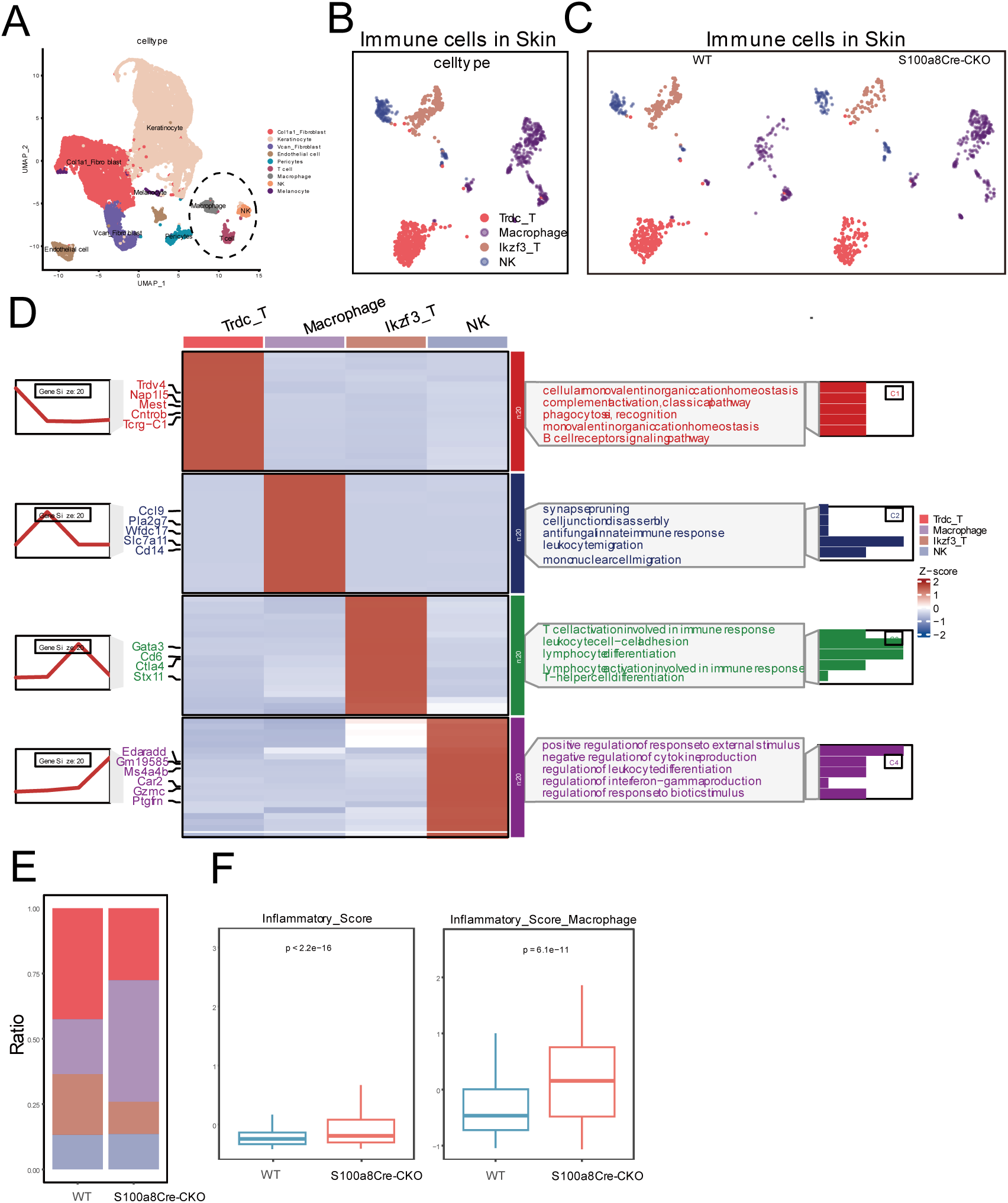
scRNA-seq analysis of skin tissues from *S100a8Cre-CKO* mice. Relying on scRNA-seq dataset, we observed a slightly increased inflammation in the skin tissue from S100a8Cre-CKO mice. Tissues were from 1 WT animal and 1 S100a8Cre-CKO animal. Related to **Figure 4** in the **Main text**. (**A-C**) Skin tissues from WT and *S100a8Cre-CKO* mice were subjected for scRNA-sequencing analysis for identifying any possible pathophysiological alterations in the *S100a8Cre-CKO* mice. Immune cells were extracted and subject for further analysis. **A**: the UMAP plot of total cells of the skin tissue; **B**: the UMAP plot of the immune cells of the skin tissue; **C**: split plots of skin immune cells. Of note, possibly due to technical problem, we failed to capture any neutrophils in the skin scRNA-seq. (**D**) The enriched pathways in each immune cell types form the skin scRNA-seq datasets. (**E**) Proportions of each immune cell types in the immune cell pool of the skin. (**F**) Scoring of the inflammatory activity for all immune cells or just macrophages.

**Figure S6:**
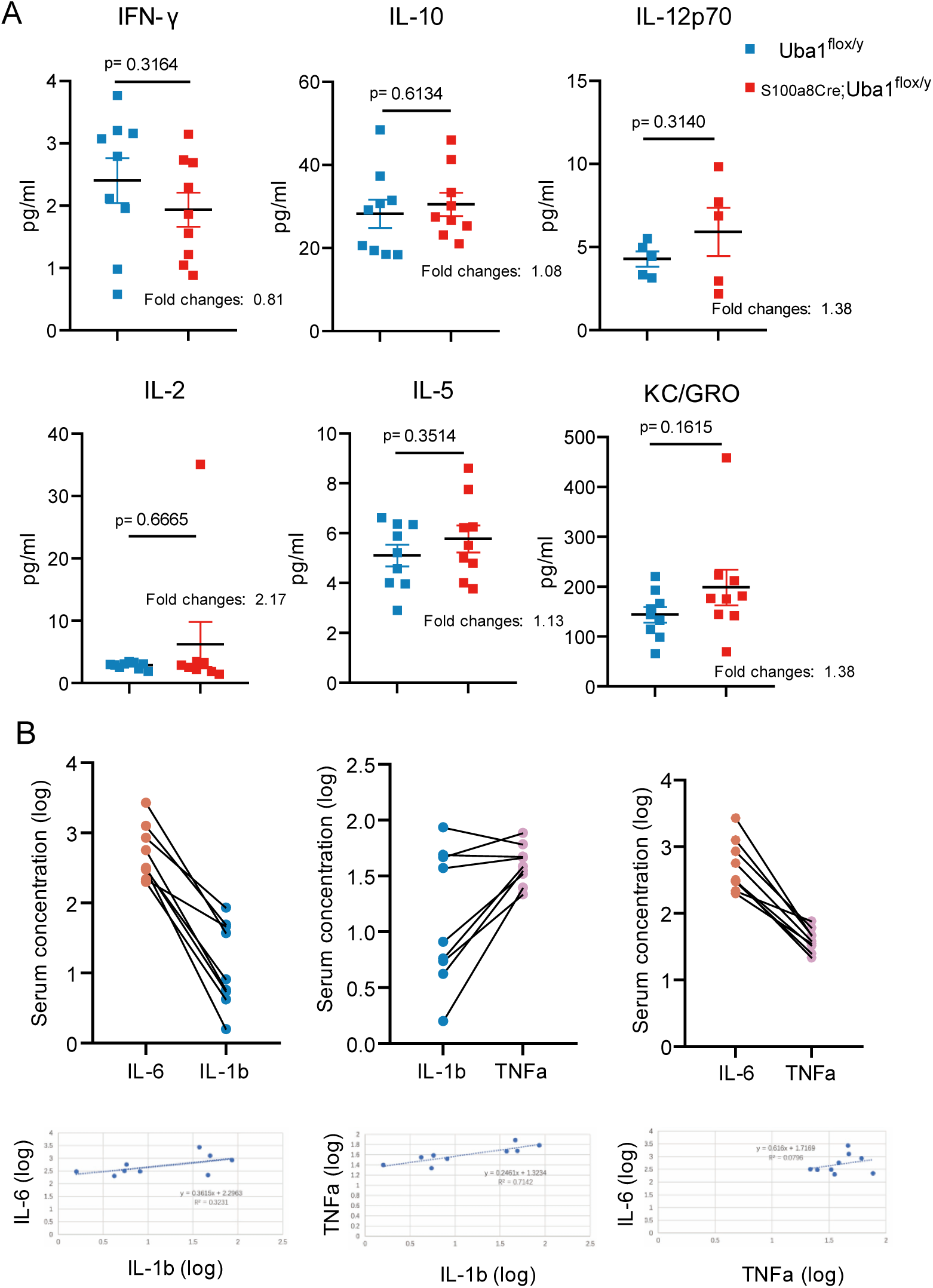
Additional ELISA results of serum cytokines from *S100a8Cre-CKO* mice. Positive correlation of IL-6, IL-1β and TNFα were observed in the serum of S100a8Cre-CKO mice. Related to **Figure 4** in the **Main text**. (**A**) Serum levels of other 6 cytokines from WT and *S100a8Cre-CKO* mice. Levels of IL-4 are undetectable in wild type and were not shown here. (**B**) Correlation analysis of pro-inflammatory cytokines IL-6, IL-1β, and TNFα from *S100a8Cre-CKO* mice.

**Figure S7:**
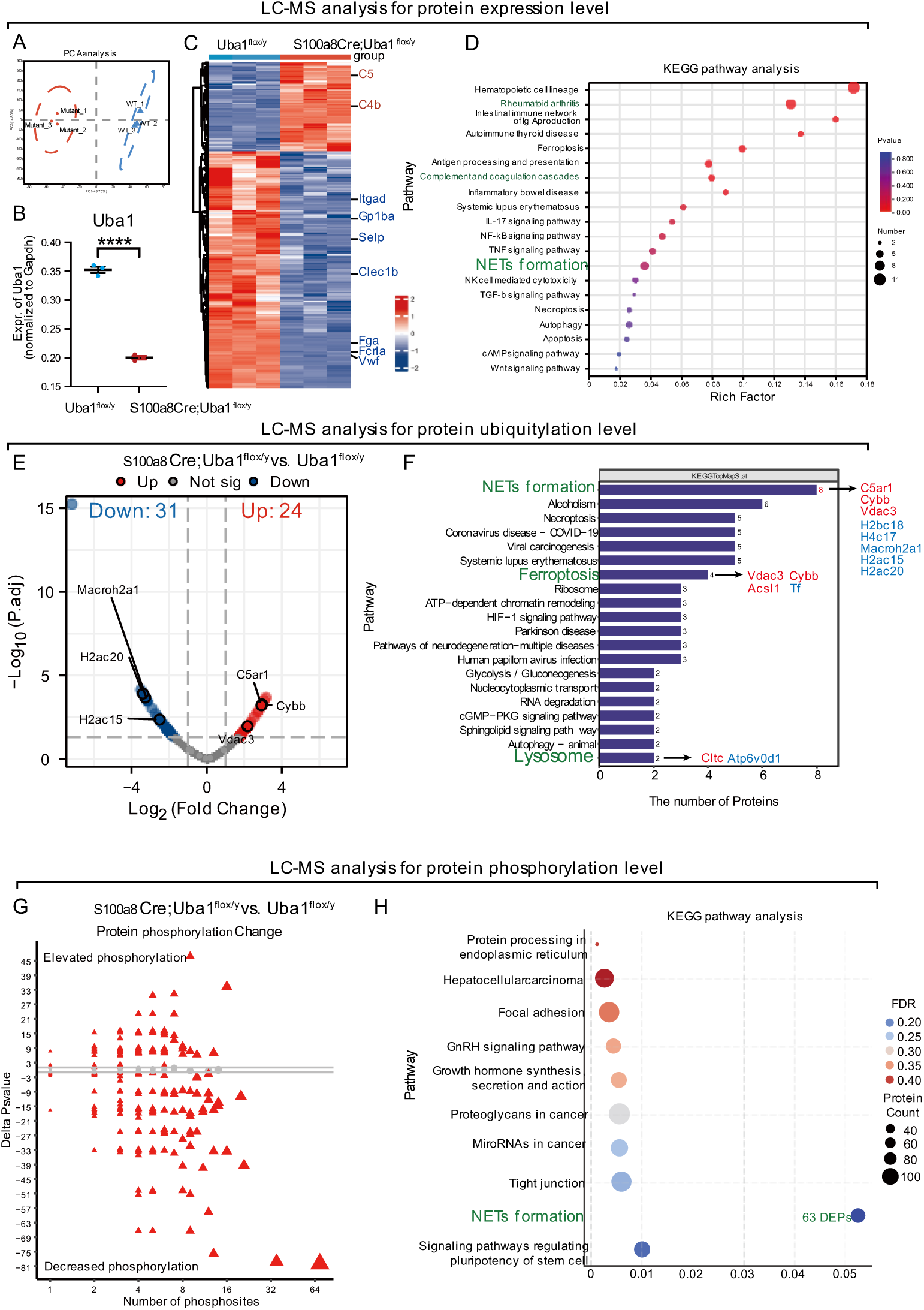
Disturbed neutrophil homeostasis in *S100a8Cre-CKO* revealed by proteomic LC-MS/MS analysis. Related to **Figure 5** in the **Main text**. BM cells (unsorted) from WT and *S100a8Cre-CKO* mice were subject for regular proteomic quantification (**A** to **D**), protein ubiquitylation measurement (**E** to **F**) and protein phosphorylation measurement (**G** to **H**). **(A)** PCA analysis of the WT (n=3 biological repeats) and *S100a8Cre-CKO* (n=3 biological repeats) BM cells used for regular protein expression level. **(B)** Expression of *Uba1* in the regular protein expression datasets were quantified (normalized to Gapdh). Fold change of Uba1 in unsorted BM cells: ∼0.57, p < 0.0001. **(C)** Heatmap of 265 different expression proteins (DEPs). In the figure, 2 up-regulated and 7 down-regulated proteins related to NET formation pathway is denoted. **(D)** Enriched pathways are highlighted when comparing *S100a8Cre* -CKO BM cells with WT BM cells. Proteins related to rheumatoid arthritis, complement cascade and NETs formation are highlighted. **(E)** Volcano plot of different expression level of ubiquitinated proteins. Proteins related to NETs formation are labeled. **(F)** Barplot of KEGG enrichment of ubiquitinated DEPs. Lysosome, NETs formation or Ferroptosis-related pathways are enriched when comparing *S100a8Cre-CKO* BM cells with WT BM cells. **(G)** Scatter plot of protein phosphorylation status values. Changes of protein phosphorylation levels (ΔPs) was calculated by summing the log2FC values of all differentially phosphorylated peptides corresponding to the same protein. ΔP value >1 or < −1 indicated that the phosphorylated peptide in the protein was up- regulated or down-regulated, represented by red triangles. −1<ΔPvalue<1 indicates that the phosphorylation in this protein has not changed, represented by grey triangles. The size of triangle means number of phosphosites. **(H)** Enriched NETs formation-related pathways are highlighted when comparing *S100a8Cre* -CKO BM cells with WT BM cells. In total 63 phosphorylated DEPs were enriched in NET formation pathway. ****, p<0.0001; n=3∼4 biological repeats.

**Figure S8:**
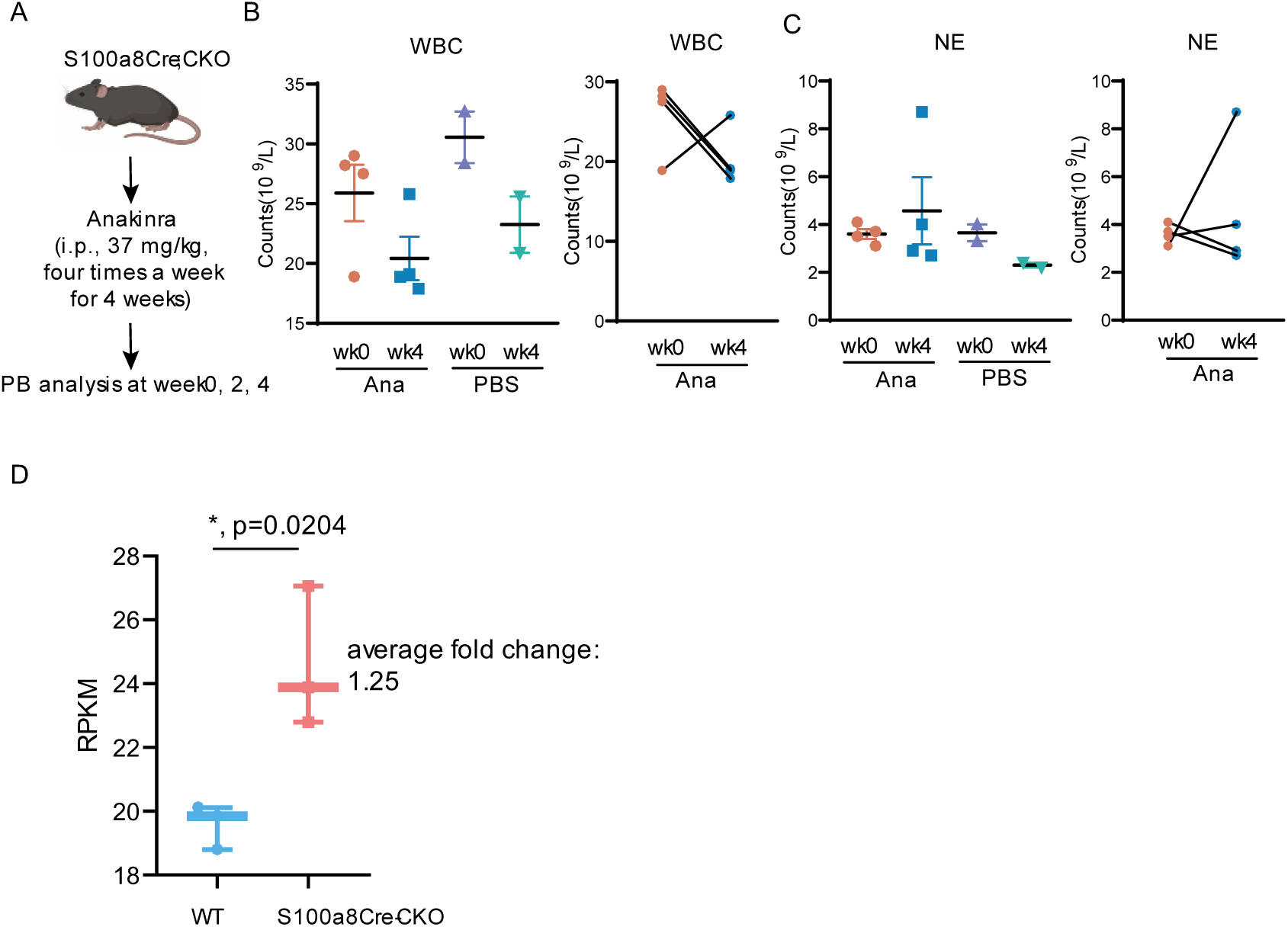
Additional results of drug treatment on the *S100a8Cre-CKO* mice. Treatment with Anakinra partially reversed abnormalities in *S100a8Cre-CKO* mice. Related to **Figure 6** in the **Main text**. (**A**) Scheme of the regime for the Anakinra treatment on *S100a8Cre* -CKO mice. (**B**) Counts of WBC at the time-points pre or post Anakinra treatment. (**C**) Counts of neutrophils at the time-points pre or post Anakinra treatment. (**D**) Expression of *Morrbid* in bone marrow cells of WT and *S100a8Cre-CKO* mice. Bulk RNA-seq dataset was normalized to RPKM for each gene. *, p<0.05; n=3∼4 biological repeats.

