## Supplementary material for "Disturbing immune homeostasis by neutrophil loss of Uba1 induces VEXAS-like autoinflammatory disease in mice": Supp. Fig1-8

### Figure S1

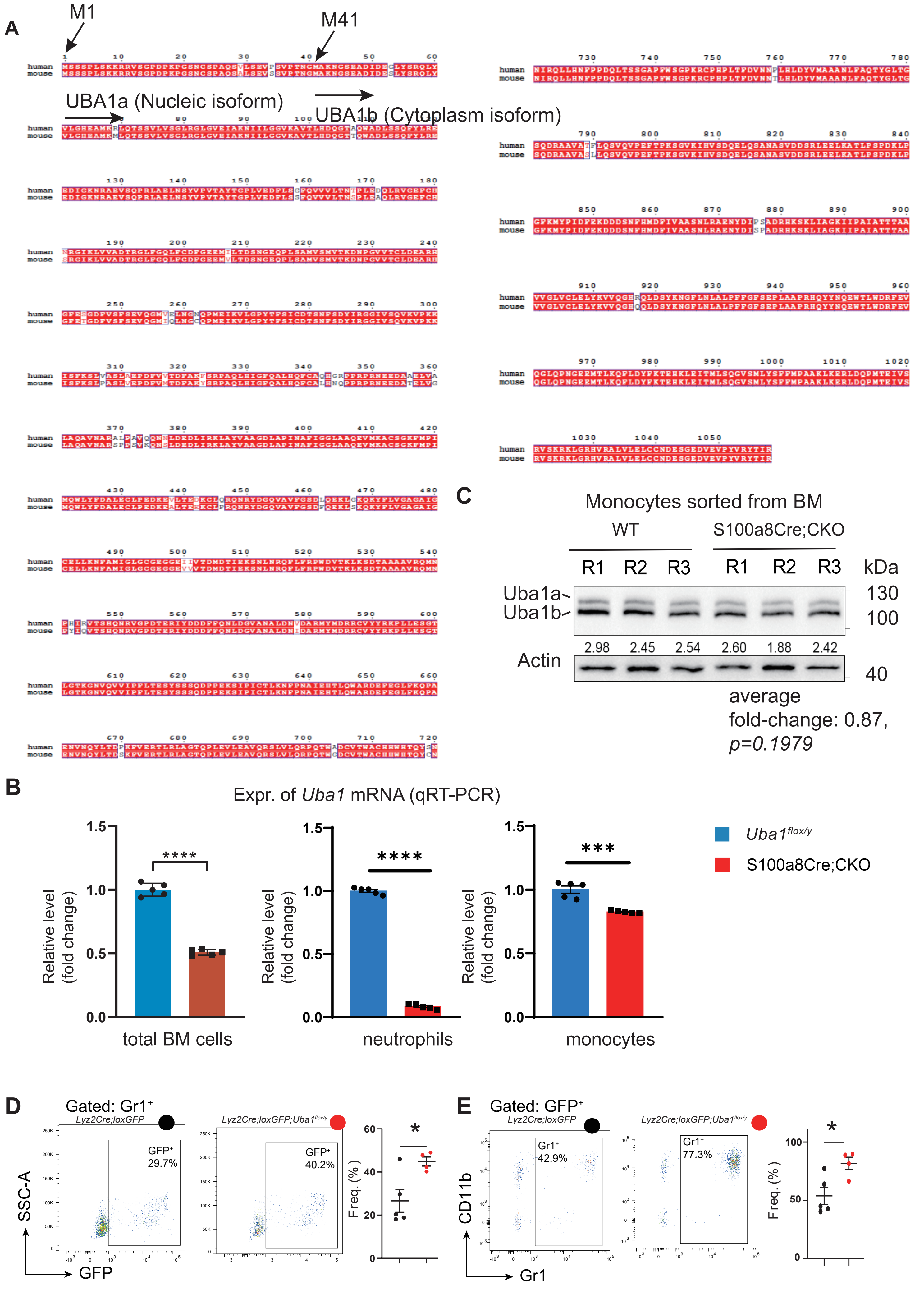

A
M1
M41
UBA1a (Nucleic isoform)
UBA1b (Cytoplasm isoform)
human
mouse
B
Expr. of Uba1 mRNA (qRT-PCR)
Relative level (fold change)
total BM cells
neutrophils
monocytes
****
****
***
Uba1flox/y
S100a8Cre;CKO
C
Monocytes sorted from BM
WT
S100a8Cre;CKO
R1 R2 R3 R1 R2 R3
kDa
Uba1a
Uba1b
130
100
2.98 2.45 2.54 2.60 1.88 2.42
Actin
40
average fold-change: 0.87, p=0.1979
D
Gated: Gr1+
Lyz2Cre;loxGFP
Lyz2Cre;loxGFP;Uba1flox/y
GFP+ 29.7%
GFP+ 40.2%
SSC-A
GFP
Freq. (%)
*
E
Gated: GFP+
Lyz2Cre;loxGFP
Lyz2Cre;loxGFP;Uba1flox/y
Gr1+ 42.9%
Gr1+ 77.3%
CD11b
Gr1
Freq. (%)
*

### Figure S2

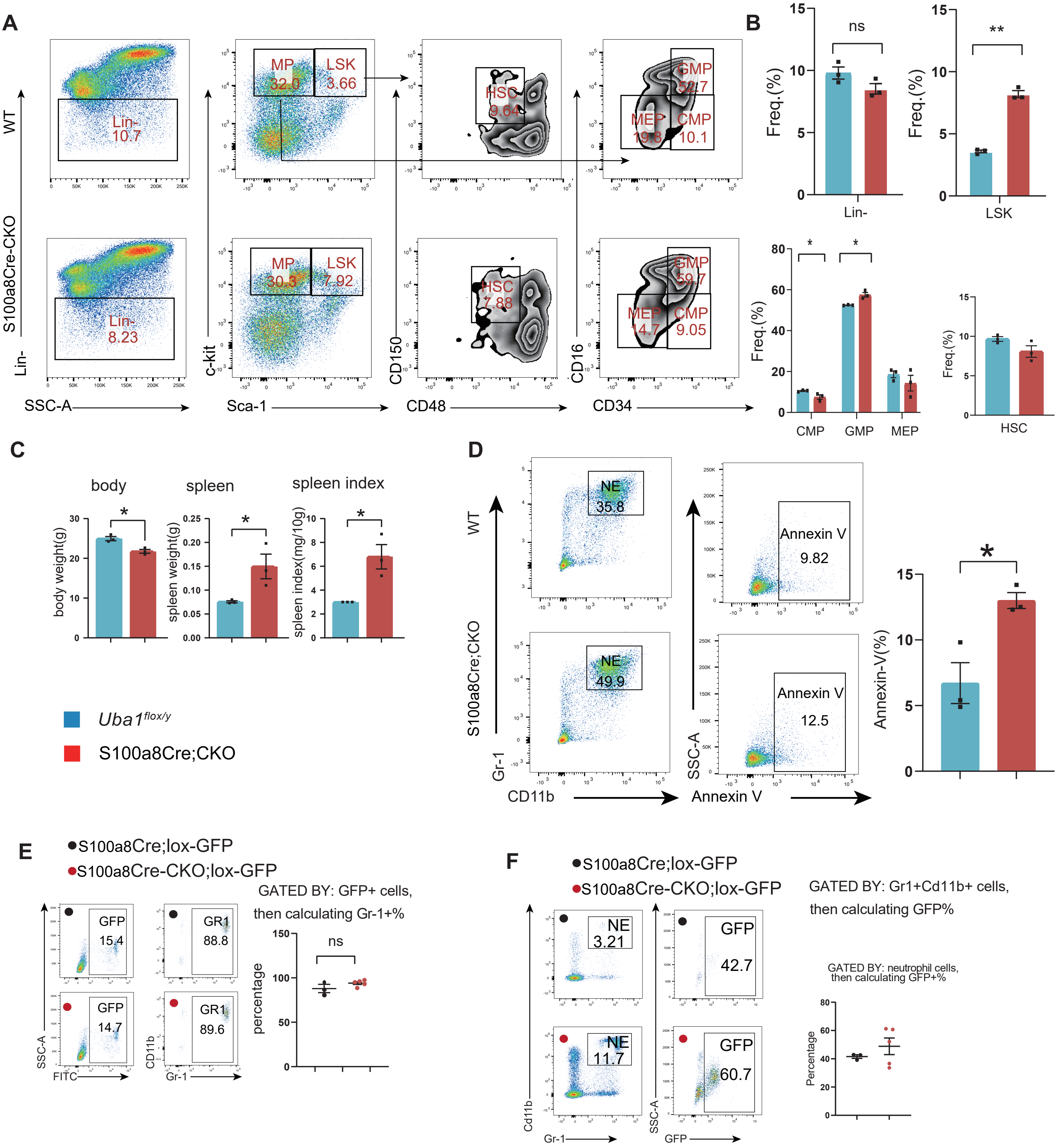

A
WT
S100a8Cre-CKO
Lin-
10.7
8.23
SSC-A
MP
32.0
LSK
3.66
30.3
7.92
c-kit
Sca-1
HSC
9.64
7.88
CD150
CD48
GMP
52.7
MEP
19.8
CMP
10.1
59.7
14.7
9.05
CD16
CD34
B
ns
**
Freq.(%)
Lin-
LSK
*
CMP
GMP
MEP
HSC
C
body
spleen
spleen index
body weight(g)
spleen weight(g)
spleen index(mg/10g)
Uba1flox/y
S100a8Cre;CKO
D
WT
S100a8Cre;CKO
NE
35.8
49.9
Annexin V
9.82
12.5
Gr-1
CD11b
SSC-A
Annexin V
Annexin-V(%)
E
S100a8Cre;lox-GFP
S100a8Cre-CKO;lox-GFP
GFP
15.4
14.7
GR1
88.8
89.6
SSC-A
FITC
CD11b
Gr-1
GATED BY: GFP+ cells,
then calculating Gr-1+%
percentage
F
S100a8Cre;lox-GFP
S100a8Cre-CKO;lox-GFP
NE
3.21
11.7
GFP
42.7
60.7
Cd11b
Gr-1
SSC-A
GFP
GATED BY: Gr1+Cd11b+ cells,
then calculating GFP%
GATED BY: neutrophil cells,
then calculating GFP+%
Percentage

**Figure S3**

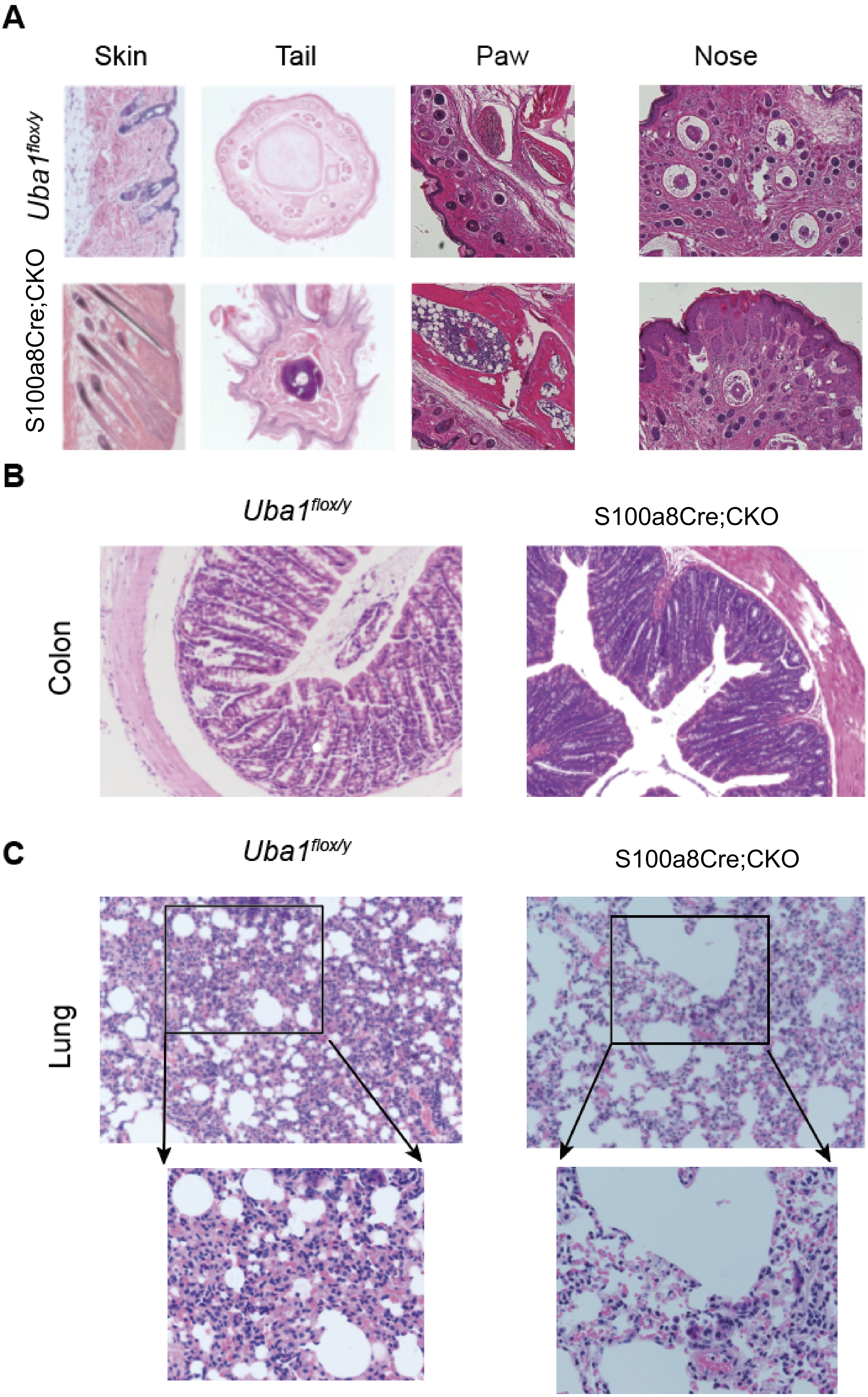
A
Skin
Tail
Paw
Nose
Uba1flox/y
S100a8Cre;CKO
B
Uba1flox/y
S100a8Cre;CKO
Colon
C
Uba1flox/y
S100a8Cre;CKO
Lung

### Figure S4

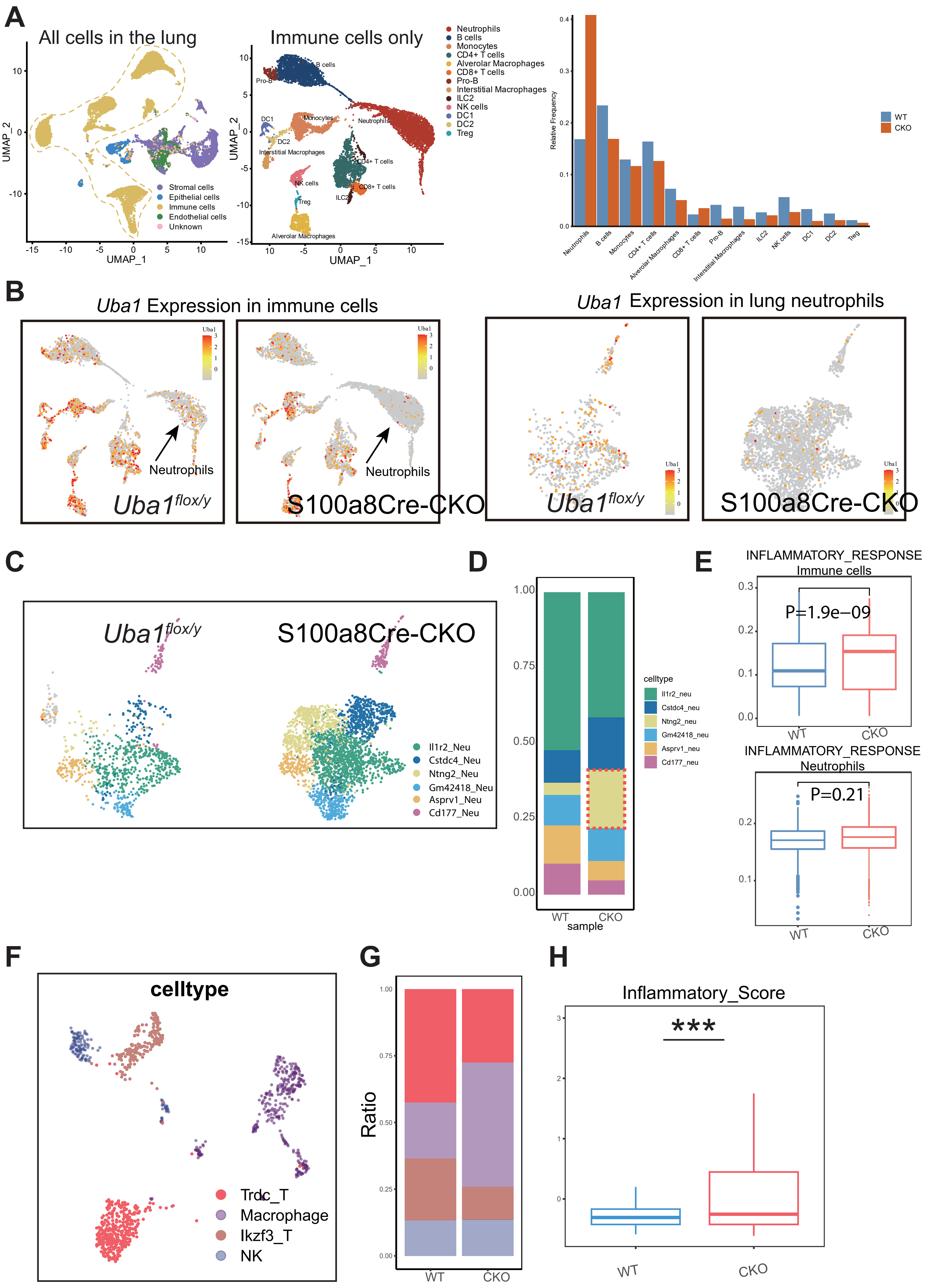

A
All cells in the lung
Immune cells only
Stromal cells
Epithelial cells
Immune cells
Endothelial cells
Unknown
UMAP_1
UMAP_2
Neutrophils
B cells
Monocytes
CD4+ T cells
Alverolar Macrophages
CD8+ T cells
Pro-B
Interstitial Macrophages
ILC2
NK cells
DC1
DC2
Treg
Relative Frequency
WT
CKO
B
Uba1 Expression in immune cells
Uba1 Expression in lung neutrophils
Neutrophils
Uba1flox/y
S100a8Cre-CKO
C
Il1r2_Neu
Cstdc4_Neu
Ntng2_Neu
Gm42418_Neu
Asprv1_Neu
Cd177_Neu
D
celltype
sample
E
INFLAMMATORY_RESPONSE
Immune cells
P=1.9e−09
Neutrophils
P=0.21
F
celltype
Trdc_T
Macrophage
Ikzf3_T
NK
G
Ratio
H
Inflammatory_Score
***

### Figure S5

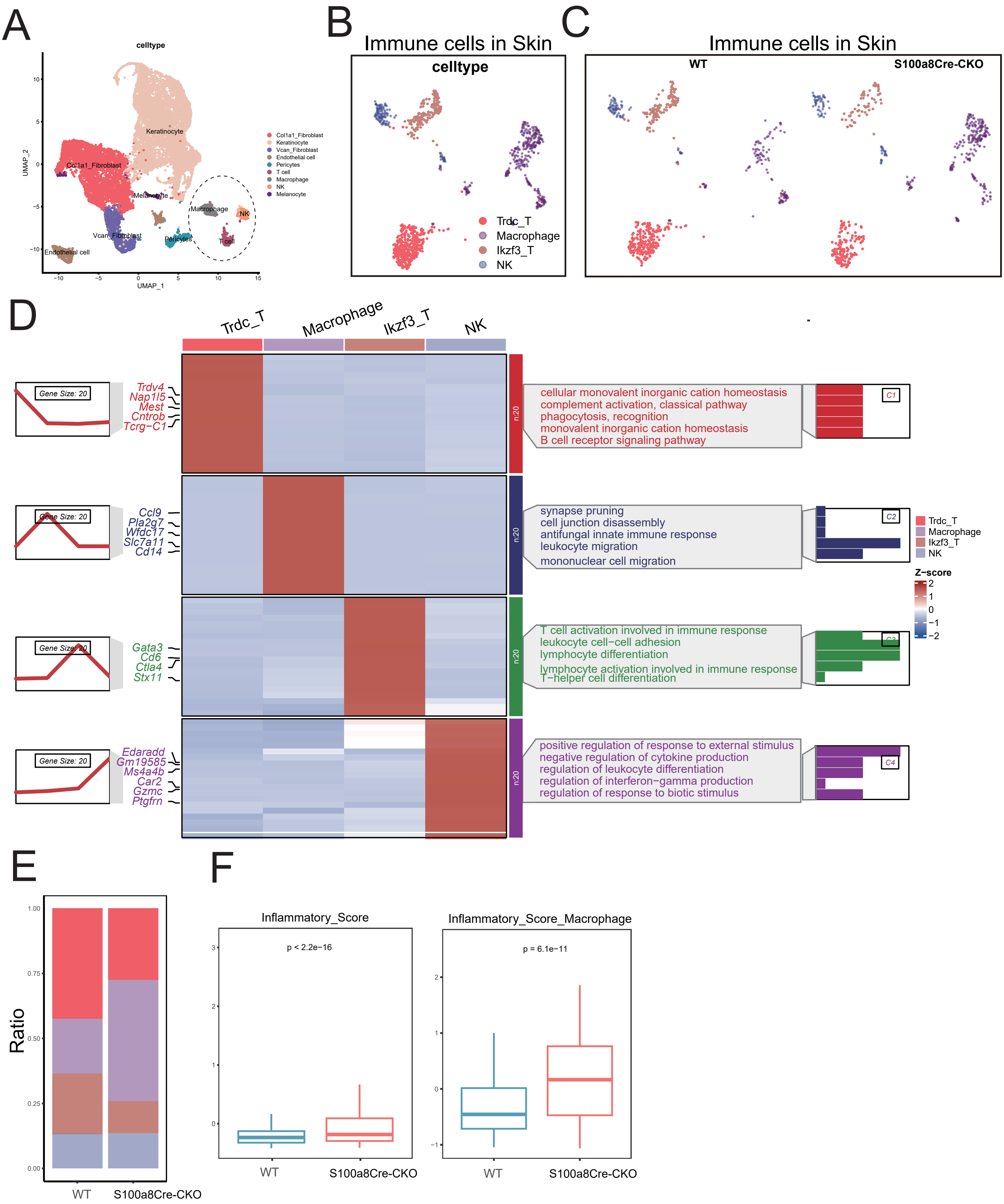

A
celltype
Keratinocyte
Col1a1_Fibroblast
Melanocyte
Macrophage
NK
T cell
Pericytes
Vcan_Fibroblast
Endothelial cell
Col1a1_Fibroblast
Keratinocyte
Vcan_Fibroblast
Endothelial cell
Pericytes
T cell
Macrophage
NK
Melanocyte
UMAP_1
UMAP_2
B
Immune cells in Skin
celltype
Trdc_T
Macrophage
Ikzf3_T
NK
C
Immune cells in Skin
WT
S100a8Cre-CKO
D
Trdc_T
Macrophage
Ikzf3_T
NK
Gene Size: 20
Trdv4
Nap1l5
Mest
Cntrob
Tcrg−C1
cellular monovalent inorganic cation homeostasis
complement activation, classical pathway
phagocytosis, recognition
monovalent inorganic cation homeostasis
B cell receptor signaling pathway
C1
Ccl9
Pla2g7
Wfdc17
Slc7a11
Cd14
synapse pruning
cell junction disassembly
antifungal innate immune response
leukocyte migration
mononuclear cell migration
C2
Gata3
Cd6
Ctla4
Stx11
T cell activation involved in immune response
leukocyte cell−cell adhesion
lymphocyte differentiation
lymphocyte activation involved in immune response
T−helper cell differentiation
Edaradd
Gm19585
Ms4a4b
Car2
Gzmc
Ptgfrn
positive regulation of response to external stimulus
negative regulation of cytokine production
regulation of leukocyte differentiation
regulation of interferon−gamma production
regulation of response to biotic stimulus
C4
n:20
Z−score
E
Ratio
1.00
0.75
0.50
0.25
0.00
WT
S100a8Cre-CKO
F
Inflammatory_Score
p < 2.2e−16
WT
S100a8Cre-CKO
Inflammatory_Score_Macrophage
p = 6.1e−11
WT
S100a8Cre-CKO

### Figure S6

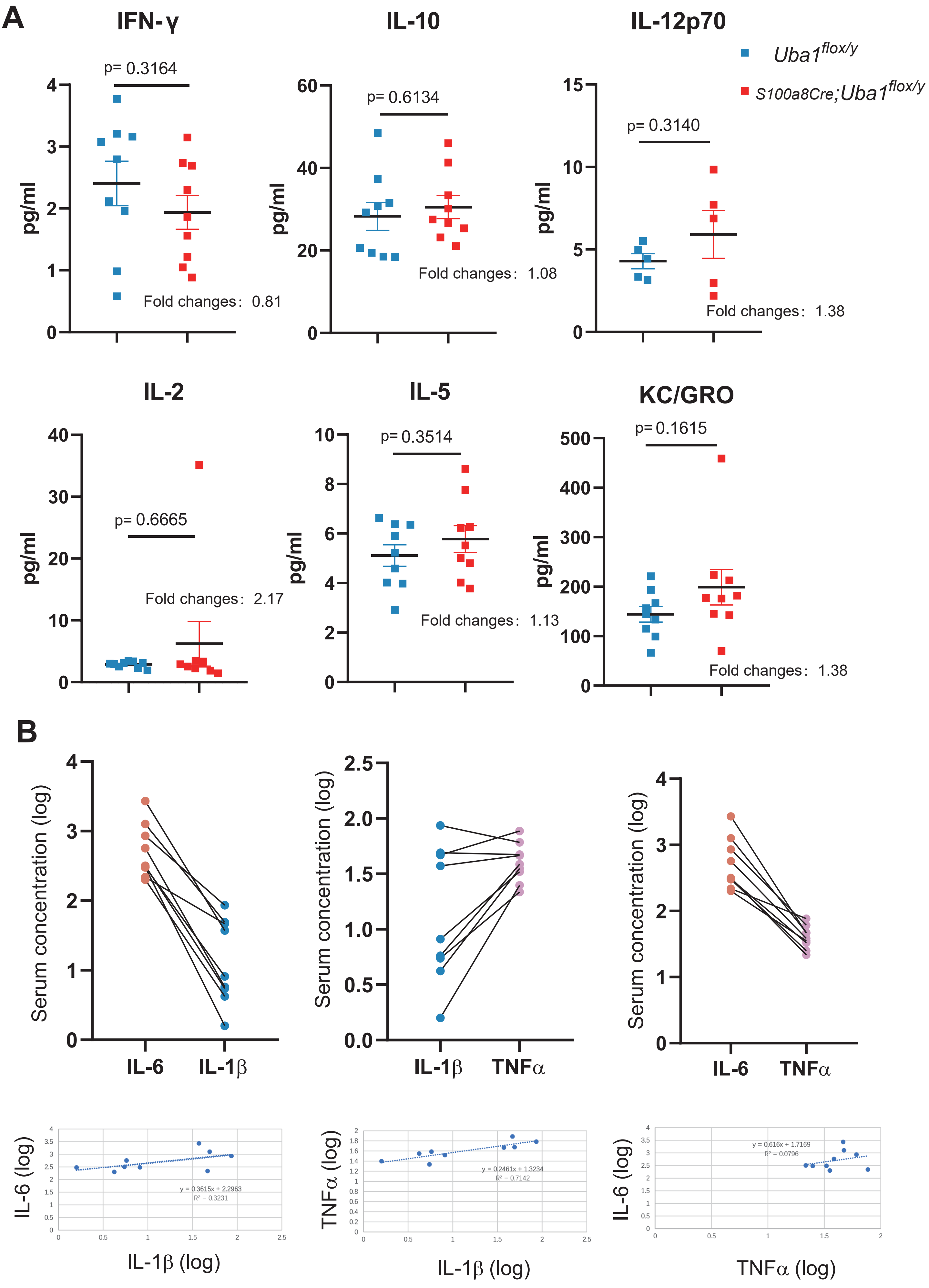

A
IFN-γ
IL-10
IL-12p70
Uba1flox/y
S100a8Cre;Uba1flox/y
pg/ml
p= 0.3164
Fold changes: 0.81
p= 0.6134
Fold changes: 1.08
p= 0.3140
Fold changes: 1.38
IL-2
IL-5
KC/GRO
p= 0.6665
Fold changes: 2.17
p= 0.3514
Fold changes: 1.13
p= 0.1615
Fold changes: 1.38
B
Serum concentration (log)
IL-6
IL-1β
TNFα
IL-6 (log)
IL-1β (log)
y = 0.3615x + 2.2963
R² = 0.3231
TNFα (log)
y = 0.2461x + 1.3234
R² = 0.7142
TNFα (log)
y = 0.616x + 1.7169
R² = 0.0796

### Figure S7

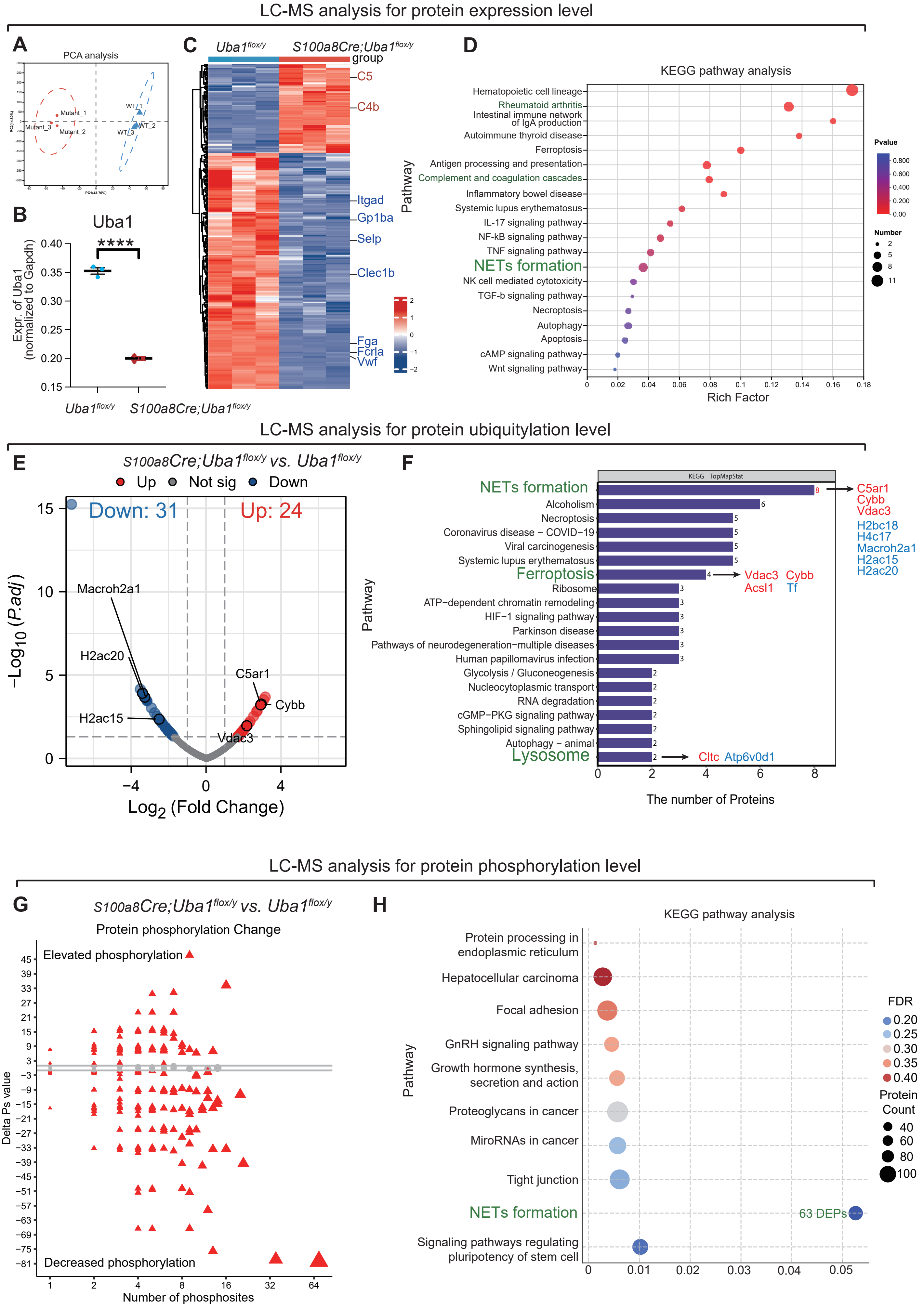
LC-MS analysis for protein expression level
A
PCA analysis
Mutant_1
Mutant_3
Mutant_2
WT_1
WT_2
WT_3
B
Uba1
****
Expr. of Uba1 (normalized to Gapdh)
0.40
0.35
0.30
0.25
0.20
0.15
Uba1flox/y
S100a8Cre;Uba1flox/y
C
Uba1flox/y
S100a8Cre;Uba1flox/y
group
C5
C4b
Itgad
Gp1ba
Selp
Clec1b
Fga
Fcrla
Vwf
2
1
0
−1
−2
Pathway
D
KEGG pathway analysis
Hematopoietic cell lineage
Rheumatoid arthritis
Intestinal immune network of IgA production
Autoimmune thyroid disease
Ferroptosis
Antigen processing and presentation
Complement and coagulation cascades
Inflammatory bowel disease
Systemic lupus erythematosus
IL-17 signaling pathway
NF-kB signaling pathway
TNF signaling pathway
NETs formation
NK cell mediated cytotoxicity
TGF-b signaling pathway
Necroptosis
Autophagy
Apoptosis
cAMP signaling pathway
Wnt signaling pathway
Pvalue
0.800
0.600
0.400
0.200
0.00
Number
2
5
8
11
0
0.02
0.04
0.06
0.08
0.1
0.12
0.14
0.16
0.18
Rich Factor
LC-MS analysis for protein ubiquitylation level
E
S100a8Cre;Uba1flox/y vs. Uba1flox/y
Up
Not sig
Down
Down: 31
Up: 24
Macroh2a1
H2ac20
H2ac15
C5ar1
Cybb
Vdac3
−Log10 (P.adj)
15
10
5
0
−4
0
4
Log2 (Fold Change)
F
KEGG TopMapStat
NETs formation
Alcoholism
Necroptosis
Coronavirus disease − COVID−19
Viral carcinogenesis
Systemic lupus erythematosus
Ferroptosis
Ribosome
ATP−dependent chromatin remodeling
HIF−1 signaling pathway
Parkinson disease
Pathways of neurodegeneration−multiple diseases
Human papillomavirus infection
Glycolysis / Gluconeogenesis
Nucleocytoplasmic transport
RNA degradation
cGMP−PKG signaling pathway
Sphingolipid signaling pathway
Autophagy − animal
Lysosome
Pathway
C5ar1
Cybb
Vdac3
H2bc18
H4c17
Macroh2a1
H2ac15
H2ac20
Vdac3
Cybb
Acsl1
Tf
Cltc
Atp6v0d1
0
2
4
6
8
The number of Proteins
LC-MS analysis for protein phosphorylation level
G
S100a8Cre;Uba1flox/y vs. Uba1flox/y
Protein phosphorylation Change
Elevated phosphorylation
Decreased phosphorylation
Delta Ps value
45
39
33
27
21
15
9
3
−3
−9
−15
−21
−27
−33
−39
−45
−51
−57
−63
−69
−75
−81
1
2
4
8
16
32
64
Number of phosphosites
H
KEGG pathway analysis
Protein processing in endoplasmic reticulum
Hepatocellular carcinoma
Focal adhesion
GnRH signaling pathway
Growth hormone synthesis, secretion and action
Proteoglycans in cancer
MiroRNAs in cancer
Tight junction
NETs formation
63 DEPs
Signaling pathways regulating pluripotency of stem cell
Pathway
FDR
0.20
0.25
0.30
0.35
0.40
Protein Count
40
60
80
100
0
0.01
0.02
0.03
0.04
0.05

### Figure S8

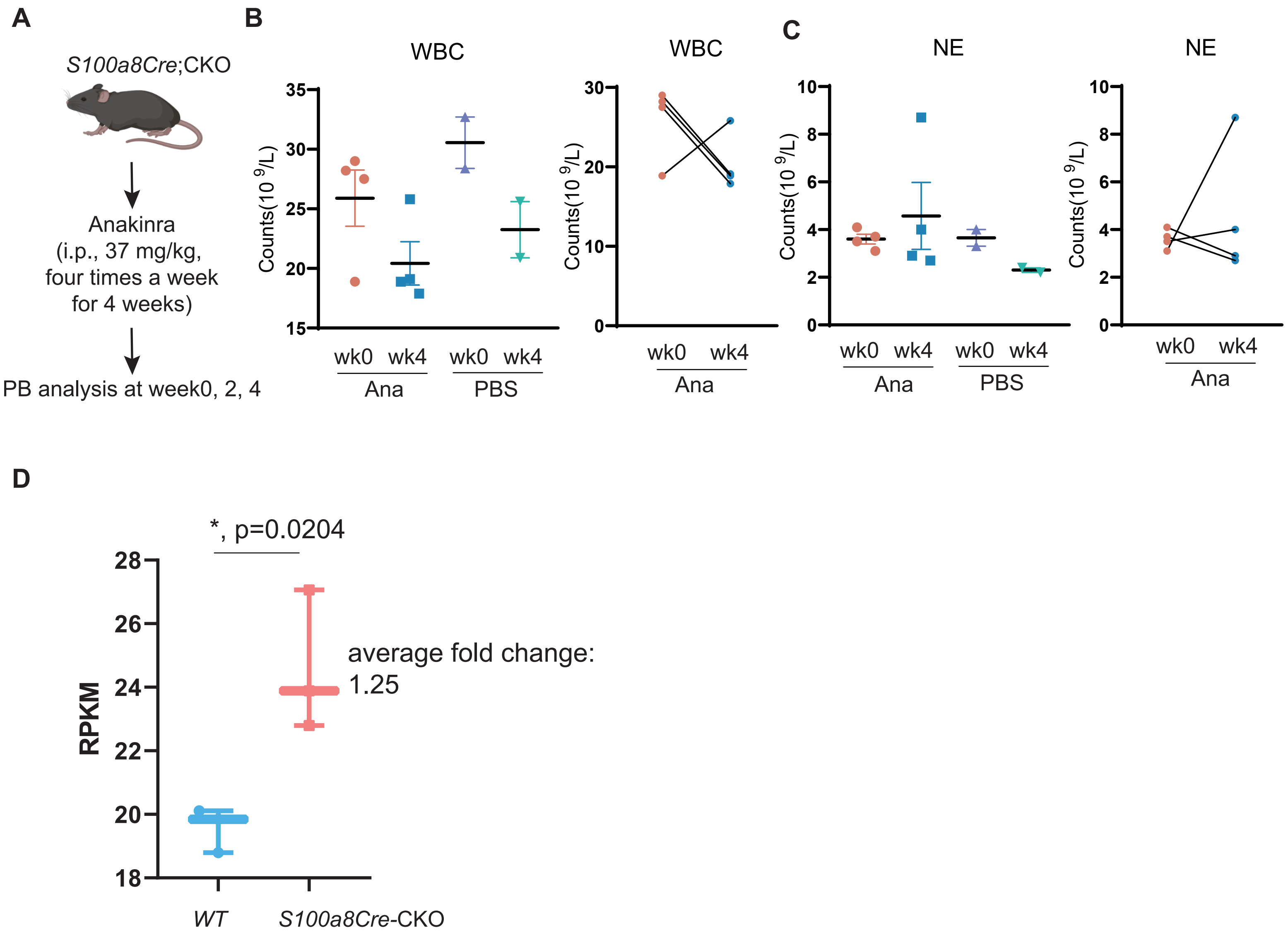

A
B
C
D
*S100a8Cre*;CKO
Anakinra
(i.p., 37 mg/kg,
four times a week
for 4 weeks)
PB analysis at week0, 2, 4
WBC
Counts(10 9/L)
wk0 wk4
Ana
PBS
NE
*, p=0.0204
average fold change:
1.25
RPKM
*WT*
*S100a8Cre*-CKO

#### Table-S1: Prevalence and penetrance of VEXAS syndrome in human population and in our CKO mice modeling

| | VEXAS Syndrome in human population | our CKO modeling in mice | Notes and References for further checking | Related Figures in our study |
|---|---|---|---|---|
| UBA1 gene location, structure and encoded proteins; | X-Chromosome, 26 exons; 1088 aa | X-Chromosome, 26 exons; 1088 aa; | ~95% identity, ~98% similarity | Figure S1; |
| Mutation Strategy | Somatic mutation, not inheritable; | Cre/Flox-based CKO strategy, cell type-dependent conditional mutations, strains are easy for maintenance by genetic breeding | Some germline mutations in UBA1 will cause XL-SMA, a neurological disease (Ref #15-16 in the Main text); | Figure 1; |
| Variants of mutation in UBA1 | VEXAS-related mutations are typically at the UBA1$^{M41}$ site, but the UBA1$^{non-M41}$ mutations were also reported (i.e. N606I, I894S, A478S, E597, S621C, P749L, P1014L) [Ref #1]. Of note, the causality of non-M41 mutations certainly require furthers functional validations. | Exon 6-8 are removed in our CKO murine modeling. Of note, removal of Exon 6-8 resulted in a shift-frame mutation and early stop codon Exon 10. By the way, Exon 6-8 in mouse and human are identical in length. | Ref #1: Sakuma, M., Blombery, P., Meggendorfer, M. et al. Novel causative variants of VEXAS in UBA1 detected through whole genome transcriptome sequencing in a large cohort of hematological malignancies. Leukemia 37, 1080–1091 (2023). https://doi.org/10.1038/s41375-023-01857-5 | Figure 1; |
| Age-related prevalence | typically for adults aged >50 | It is also age-dependent for the S100a8Cre; CKO model. About 10% S100a8Cre; CKO mice show up phenotypes around 1-2 month old. Most of the mutant mice show up phenotype starting at 4-6 month old. | Age and VAF are correlated; | Figure 4; |
| Gender-related prevalence | VEXAS syndrome is typically identified in adult males; There is a very rare occurrence of VEXAS syndrome in adult female. | Both male and female CKO mutants could be produced by the Cre/Flox CKO strategy. Importantly, both male and female S100a8Cre; CKO manifested similar VEXAS-like symptoms in our study. | We observed similar VEXAS-like phenotypes in female S100a8Cre;CKO | N.A. |
| *VAF dependency | The authors believe VEXAS is not only an age-dependent disease but also a VAF-dependent disease. Further clinical and experiment studies are demanded to validate this. | According to our experimental data, we approximated the VAF of Vav1Cre, R26CreErt2 and Mx1Cre-based CKO in HSCs is close to 100%; we approximated the VAF of Cd4Cre, Cd19Cre and Pf4Cre based CKO in the specific cell types is close to 50%; we approximated the VAF of Lyz2Cre-based CKO in the monocytes is close to 50% (according to loxGFP tracing data); we approximated the VAF of S100a8Cre-based CKO in the monocytes is close to 70% (according to data from different measurements)**; | *VAF: variant allele frequency; VAF is a very important measurement of somatic mutations for entities including leukemia, clonal hematopoiesis and VEXAS. **We used the depletion efficiency (y) value of Uba1 to approximate the VAF of Uba1 in certain cell types. | N.A. |

#### Table-S2: Cell-type-dependent tolerance and pathogenicity in pathology of VEXAS Syndrome patients and our 9 CKO models

| | Tolerance and pathogenicity in human VEXAS syndrome patients [Ref#2] | Tolerance and pathogenicity in our CKO mice modeling | Notes and References for further checking | Related Figures in our study |
|---|---|---|---|---|
| HSC* | yes, tolerated according to Figure 1 of Ref #2; Toxicity (pathogenicity) of HSC* is imaginable since it is on the top of hematopoietic system but further experimental, perturbation and functional studies are demanded for the details, in addition the recent availability of scRNA-seq datasets of VEXAS syndrome patients) | not tolerated for null-mutation of Uba1, resulted in death of HSC*, | * means mutant cells carrying UBA1 mutations (somatic mutations in human samples; a null mutation in our CKO models); Ref#2: most of the cell type-related pathology information is based on NEJM-2020, the first report of VEXAS (PMID: 33108101); The authors are planning a meta-study to cover all of the available clinical observations. | Figure 2; |
| GMP* | yes, tolerated according to Figure 1 of Ref #2. But the authors believe further functional experiments are needed to demonstrate its pathogenicity and the detail mechanisms; | N.A. We don't have direct data for this cell type but please check our speculations on the right column; | Based on our data from HSC-CKO models, we speculate the null mutation of Uba1 in GMP also cause death of GMP; | N.A. |
| MEP* | yes, tolerated according to Figure 1 of Ref #2; but the authors believe further functional experiments need to be conducted to demonstrate its pathogenicity; | N.A. We don't have direct data for this cell type; but please check our speculations on the right column; | Based on our data from HSC-CKO models, we speculate the null mutation of Uba1 in MEP will cause death of MEP; | N.A. |
| Erythroid blast* | yes, tolerated according to Figure 1 of Ref #2. | N.A. We don't have direct data for this cell type; but please check our speculations on the right column; | Based on our limited data from HSC-CKO models, we speculate the null mutation of Uba1 in Erythroid blast will cause death of Erythroid blast; | N.A. |
| Megakaryocyte* | yes, tolerated according to Figure 1 of Ref #2. | not tolerated for null-mutation, resulted in death of Megakaryocytes*, | Based on Pf4Cre; CKO, we demonstrated that megakaryocyte is not tolerated to null mutation of Uba1; | Figure 3; |
| LymP* (Lymphoid Progenitor) | yes, tolerated according to Figure 1 of Ref #2; but the authors believe further functional experiments is required to demonstrate its toxicity; | N.A. We don't have direct data for this cell type; but please check our speculations on the right column; | Based on our data from HSC-CKO models, we speculate the null mutation of Uba1 in LymP also cause death of mutant LymP; | N.A. |
| B cells* | not tolerated; and may induce dysfunction of adaptive immunity | not tolerated for null-mutation, resulted in death of mutant B cells; | Based on Cd19Cre; CKO, we demonstrated that B cell is NOT tolerated to null mutation of Uba1 | Figure 3; |
| T cells* | not tolerated; and may induce dysfunction of adaptive immunity | not tolerated for null-mutation, resulted in death of mutant T cells; | Based on Cd4Cre; CKO, we demonstrated that T cell is NOT tolerate to null mutation of [illegible] | Figure 3; |
| Monocytes* | yes, tolerated; and probably toxic; further functional experiments are needed to demonstrate its toxicities; | Based on PB cell counting, flow cytometry and lox-GFP tracing, we think mutant monocytes are tolerated to null mutation of Uba1 | Based on Lyz2Cre; CKO and Cx3cr1Cre; CKO, we demonstrated that monocytes/macrophages are tolerated to null mutation of Uba1 | Figure 3; |
| Neutrophils* | yes, tolerated and probably toxic; further functional experiments are needed to demonstrate its toxicities; | Based on PB cell counting, flow cytometry, cBMT tracing, lox-GFP tracing, BrdU and scRNA-seq, we think mutant neutrophils are tolerated to null mutation of Uba1; Of note, based Annexin-V flow cytometry, mutant neutrophils manifest higher apoptosis profile, thus further experimentation is required to clarify the relationship between tolerance, apoptosis and necroptosis. | Based on S100a8Cre; CKO, we demonstrated that mutant neutrophils are tolerated to null mutation of Uba1. | Figure 4-6; Figure S2; |

**Table-S3: Hematological and other tissue symptoms in human VEXAS Syndrome and in VEXAS-like S100a8Cre;CKO mice**

| | Symptoms in human VEXAS Syndrome [Ref#2] | Symptoms in our S100a8Cre;CKO mice | Notes and References for further checking | Related Figures in our study |
|---|---|---|---|---|
| **Part_1: Clinical features in Blood and Bone marrow** | | | | |
| WBC counts | average value is within the normal range according to Figure S3 of Ref#2 | Significantly increased | Ref#2: the symptoms of VEXAS syndrome are adapted from NEJM-2020, the first report of VEXAS (PMID: 33108101); | Figure 4D and E; |
| RBC counts | N.A. but subtle anemia is speculated according to Figure S3 of Ref#2 | Slightly decreased | | data was recorded but not shown in the Main text; |
| Platelet counts | Slightly decreased according to Figure S3 of Ref#2 | Appear normal | | data was recorded but not shown in the Main text; |
| MCV | Significant increased according to Figure S3 of Ref#2 | Significant increased | | Figure 4D and E; |
| Neutrophil counts | average value is within the normal range according to Figure S3 of Ref#2 | Significant increased | | Figure 4D and E; |
| Monocyte counts | average value is below the normal range according to Figure S3 of Ref#2 | Appear normal | | data was recorded but not shown in the Main text; |
| Neutrophil % | average value is within the normal range according to Figure S3 of Ref#2 | Significant increased | | Figure 4D and E; |
| T-cell counts | average value is below the normal range according to Figure S3 of Ref#2 | appear normal in the S100a8Cre;CKO | | Figure 4D and E; |
| B-cell counts | average value is within the normal range according to Figure S3 of Ref#2 | Significantly decreased in the S100a8Cre;CKO | | Figure 4D and E; |
| Serology of cytokines | CRP is dramatically increased while IL-8, IP-10 and INFg are slightly increased according to Figure S3 of Ref#2 | Proinflammatory cytokines IL-6, IL-1 and TNFa are significantly increased; | | Figure 4H and Figure S6; |
| Histology of blood cells | Vacuoles are observable in the BM and PB of syndrome patients; Of note, the frequency and penetrance of vacuoles is only about 14% according to the Figure Table S14. | Vacuoles are observable in the PB of S100a8Cre; CKO; Of note, the frequency penetrance of vacuoles is only about 10% according to our own data. | It is worthwhile to note that only about 10~14% myeloid cells manifest vacuoles in both human clinic samples and our VEXAS-like murine models; | Figure 4G; |
| **Part_2: Additional clinical features** | | | | |
| Relapsed Polychondritis (RP) | RP is pretty frequent associated with VEXAS syndrome in the patients; The prevalence of RP in VEXAS is about 60% according to Ref#2; | We observed only flare noses but the chondritis is undetectable in our S100a8Cre; CKO mice raised in SPF facility | | Figure 4A; |
| Abnormalities in Lung | Pulmonary infiltration (penetrance: 72%) | Lung histology appears normal but the scRNA-seq analysis suggest there is inflammation. | | Figure 4A; Figure S3; Figure S4; |
| Abnormalities in Skin | dermatitis and vasculitis | hair loss; dry skin; | | Figure 4A; Figure S3; Figure S5; |
| Tail | N.A. | dry skin in the tail; kink tails | | Figure 4A; Figure S3; |
| Fingers and toes | N.A. | Swollen toes (penetrance: ~ 30%); pigmentation* in the toes (penetrance 100%) | *Abnormal pigmentation on fingers and toes are reported in clinic according to the authors' knowledge; | Figure 4A; |
| **Part_3: Additional *in vitro* experimental test** | | | | |
| NETs for neutrophils* | Slightly higher NETs according to Figure S10 of Ref#2 (p=0.02) | We did not perform a experimental test. Based on the scoring of NETs by scRNA-seq and proteomic analysis, we indicated that NETs is activated in S100a8Cre;CKO; | | Figure 4D and E; Figure S7; |
| Phagocytosis for neutrophils | Slightly higher phagocytosis according to Figure S10 of Ref#2 (p=0.73) | Significant increased according to our phagocytosis experimental test | | Figure 6A and B; |

#### Table-S4: Current treatment studies in human VEXAS syndrome and in VEXAS-like S100a8Cre;CKO mice

| | Treatments in human VEXAS Syndrome patients | Treatments in our S100a8Cre;CKO mice | Notes and References for further checking | Related Figures in our study |
|---|---|---|---|---|
| **Part_1: Anti-inflammation strategies** | | | | |
| IL1-IL1R1 pathway inhibitors | About 50~80% CR according to Ref #3; | Anakinra and Canakinumab were tested with limited tries of regimes in our study. Only partial rescue were observed based on the current regimes; | Ref #3: Boyadzhieva Z, Ruffer N, Kötter I, Krusche M. How to treat VEXAS syndrome: a systematic review on effectiveness and safety of current treatment strategies. Rheumatology (Oxford). 2023 Nov 2;62(11):3518-3525. doi: 10.1093/rheumatology/kead240. PMID: 37233149 | Figure 6C-D and Figure S8A-C; Further regime tries are demanded to clarify the treatment effect of Anakinra and Canakinumab in our S100a8Cre;CKO. |
| Targeting Morrbid | N.A. But numerous reports, including our previous studies, suggest that increased expression of Morrbid is associated with leukemia and Hypereosinophilic Syndrome (HS). | Partially rescued and with good response especially for the serum cytokines | Ref #: 48-51 in the Main text. | Figure 6E-F; |
| JAK2 inhibitors | 2 out of the 3 cases in short-term treatment show up amelioration according to Ref #4; | N.A. | Ref #4: Salehi T, Callisto A, Beecher MB, Hissaria P. Tofacitinib as a biologic response modifier in VEXAS syndrome: A case series. Int J Rheum Dis. 2023 Nov;26(11):2340-2343. doi: 10.1111/1756-185X.14785. Epub 2023 Jun 19. PMID: 37337622. | N.A. It would be interesting to test the treatment with Jak2 inhibitors in S100a8Cre;CKO in the future study; |
| **Part_2: Bone marrow transplantation strategies** | | | | |
| Bone marrow transplantation treatment | Couple of clinical studies suggest bone marrow transplantation amelioted the symptoms of VEXAS syndrome. See Ref #5-6 | N.A. | Ref #5: Bone Marrow Transplant. 2022 Nov;57(11):1642-1648. doi: 10.1038/s41409-022-01774-8. Epub 2022 Aug 8. PMID: 35941354. Ref #6: Blood Adv. 2024 Mar 26;8(6):1444-1448. doi: 10.1182/bloodadvances.2023012478. PMID: 38330178; PMCID: PMC10955646. | N.A. It would be interesting to test BMT assays using S100a8Cre;CKO as recipients in future study; |
